# Tyrosine phosphorylation-dependent localization of TmaR, a novel *E. coli* polar protein that controls activity of the major sugar regulator by polar sequestration

**DOI:** 10.1101/2020.08.01.232603

**Authors:** Tamar Szoke, Nitsan Albocher, Sutharsan Govindarajan, Anat Nussbaum-Shochat, Orna Amster-Choder

## Abstract

The poles of *E. coli* cells are emerging as hubs for major sensory systems, but the polar determinants that allocate their components to the pole are largely unknown. Here, we describe the discovery of a novel protein, TmaR, which localizes to the *E. coli* cell pole when phosphorylated on a tyrosine residue. TmaR is shown here to control the subcellular localization of the general PTS protein Enzyme I (EI) by preventing it from exerting its activity by binding and polar sequestration, thus regulating sugar uptake and metabolism. Depletion or overexpression of TmaR results in EI release from the pole or enhanced recruitment to the pole, which leads to increasing or decreasing the rate of sugar consumption, respectively. Notably phosphorylation of TmaR is required to release EI and enable its activity. Like TmaR, the ability of EI to be recruited to the pole depends on phosphorylation of one of its tyrosines. In addition to hyperactivity in sugar consumption, the absence of TmaR also leads to detrimental effects on the ability of cells to survive in mild acidic conditions. Our results argue that this survival defect, which is sugar- and EI-dependent, reflects the difficulty of cells lacking TmaR to enter stationary phase. Our study identifies TmaR as the first *E. coli* protein reported to localize in a tyrosine-dependent manner and to control the activity of other proteins by their polar sequestration and release.

**SIGNIFICANCE:** In recent years, we have learnt that bacterial cells have intricate spatial organization despite the lack of membrane-bounded organelles. The endcaps of rod-shaped bacteria, termed poles, are emerging as hubs for sensing and responding, but the underlying mechanisms for positioning macromolecules there are largely unknown. We discovered a novel protein, TmaR, whose polar localization depends on a phospho-tyrosine modification. We show that TmaR controls the activity of EI, the major regulator of sugar metabolism in most bacteria, by polar sequestration and release. Notably, TmaR is essential for survival in conditions that *E. coli* often encounters in nature. Hence, TmaR is a key regulator that connects tyrosine phosphorylation, spatial regulation, sugar metabolism and survival in bacteria and the first protein reported to recruit proteins to the *E. coli* cell poles.

## INTRODUCTION

The central dogma describes the flow of genetic information from DNA to RNA to protein. However, for this process to be successful, the final product - the protein - needs to be in the right place, where it is required, and at the right time. The consequences of mislocalization can be harmful to any cell type, let alone to the unicellular bacterial cell, whose survival depends on fast and efficient response to environmental changes. Hence, protein localization is an important post-translational regulatory step. Thus far, most examples of protein targeting were reported in eukaryotic cells, usually in the context of transport from one organelle to another (*1*). In recent years, it became evident that localization of both proteins and RNAs to specific subcellular domains occurs also in prokaryotic cells, and is vital for many cellular processes (*2–5*). However, the mechanisms underlying macromolecules targeting to specific subcellular domains in bacterial cells, with the exception of membrane and cell division proteins, remain largely unknown.

The bacterial cell poles are emerging as important domains that accommodate protein and RNA assemblies (*5, 6*). Pole-localized proteins are involved in a wide range of cellular functions, including motility, regulation of cell cycle, metabolism, differentiation, pathogenesis and secretion (*7*). Various proteins are kept at the *E. coli* cell pole as inactive till they are needed, e.g., MurG (*8*), and FtsZ (*9*). On the other hand, most proteins of the chemotaxis complex act at the pole, except for CheY that is activated at the pole, but leaves to exert its effect on the flagella rotors located around the cell circumference (*10*). The phosphotransferase system (PTS), which controls sugar utilization and metabolism in most bacteria, provides another example for regulation via polar cluster formation. Execution of the PTS functions depends on a phosphorylation cascade that initiates with EI and HPr - the general PTS proteins - that deliver the phosphate to the PTS sugar permeases, which import and phosphorylate the incoming sugars (*11*). The PTS proteins also exert different effects on non-PTS proteins depending on their phosphorylation on histidine residues, thus modulating the hierarchy in sugar utilization (*11*). The PTS-imported sugars enter glycolysis, whose product, phosphoenolpyruvate (PEP), phosphorylates EI, making EI an important link between glycolysis and sugar uptake (*12*). We have previously shown that the general PTS proteins localize to the *E. coli* cell poles (*13*), although their localization depends on yet unknown factors (*14*), that during growth, EI polar clusters form stochastically from pre-existing dispersed molecules, and that EI clustering negatively correlates with EI function (*15*). Still, conclusive proof for polar localization as an inhibitory mechanism of EI function is lacking and the identity of the factor that captures EI at the pole remained unknown.

Polar clusters offer additional benefits, such as communication between signal transduction systems in order to generate an optimal response, e.g., the chemotaxis and the PTS system (*16*), or the establishment of cellular asymmetry to coordinate developmental programs with cell-cycle progression (*7*). Polar proteins that recruit other proteins to the poles, thus regulating cell cycle progression, were discovered in some bacteria, e.g., DivIVA in *Bacillus subtilis*, PopZ and TipN in *Caulobacter crescentus* and HubP in *Vibrio cholerae* (*7*). However, no pole-localizing protein has been identified in *E. coli* thus far.

In some bacteria, e.g., *C. crescentus*, specific localization of proteins is linked to their phosphorylation or to the phosphorylation of factors regulating them (*17, 18*). In most cases, these proteins are members of the two-component systems, which mediate sensing and regulation by phosphorylation on histidine and aspartic acid residues (*19*). These phosphorylation events have been considered as most prevalent in bacteria. Only in recent years, the combination of improved methodologies revealed numerous previously unknown Ser/Thr/Tyr phosphorylation sites, once thought to be hallmarks of phosphorylation in eukaryotes, in bacterial and archaeal proteins. The degree to which these putative sites are phosphorylated is still unclear and proofs for their roles and importance *in vivo* are just beginning to emerge. Also, the linkage between phosphorylation on Ser/Thr/TyR and localization of the phosphorylated bacterial proteins remained unknown.

In this study, we show that a previously uncharacterized *E. coli* protein, YeeX, which is prevalent among Gram-negative bacteria, clusters at the pole in a tyrosine phosphorylation-dependent manner and recruits the major sugar utilization regulator EI. We, therefore, renamed this protein TmaR for <u>t</u>argeting of sugar <u>m</u>etabolism-<u>a</u>ssociated <u>r</u>egulator. TmaR and EI are shown to physically interact and to co-localize. TmaR is necessary for EI polar clustering, but the opposite is not true. Only phosphorylated TmaR can release EI from the poles, since the diffuse non-phosphorylated TmaR binds to EI quite irreversibly. Notably, tyrosine phosphorylation of EI is also requires for its polar localization. We further show that TmaR-mediated EI clustering inversely correlates with EI-mediated sugar uptake, implying that the polar clusters serve as a reservoir for ready-to-act EI molecules. Cells lacking TmaR have detrimental effects on cell survival in an EI concentration- and sugar concentration-dependent manner when challenged with mildly acidic conditions, typical to various *E. coli* habitats. Taken together, our study identifies TmaR as a spatial regulator of sugar metabolism and bacterial survival and as the first reported *E. coli* pole-localizer.

## RESULTS

### TmaR assembles into polar clusters in a tyrosine phosphorylation-dependent manner

To identify yet unexplored polar proteins, we screened by fluorescence microscopy the library constructed by Xie and coworkers of 1018 *E. coli* genes tagged with YFP (*20*). This library allows the observation of C-YFP-tagged proteins expressed from their corresponding loci and native promoters in the chromosome. We thus identified TmaR (previously designated YeeX), a 109 residue-long protein of unknown function predicted to form a coiled-coil structure (https://zhanglab.ccmb.med.umich.edu/QUARK/), which is conserved among Gram-negative bacteria (**see SI Appendix, Fig. S1)**. The TmaR-YFP protein formed clusters that localized mainly near one pole of the rod-shaped *E. coli* cells (**Fig. 1A**). Measurements of the YFP intensity along the long axis of clusters-containing cells demonstrated that the vast majority of these cells had a peak in one pole (**Fig. 1B**), indicating that they have a TmaR-YFP cluster localizing to one of the poles. To verify that this localization is not due to the fluorescent tag and is not affected by the site of fusion, we monitored the localization of a TmaR that was N terminally tagged with monomeric YFP and expressed from its native chromosomal locus and promoter. The results (see SI Appendix, **Fig. S2**) show that TmaR forms polar clusters regardless of the type of tag and of which terminus is tagged.

**Figure 1.**
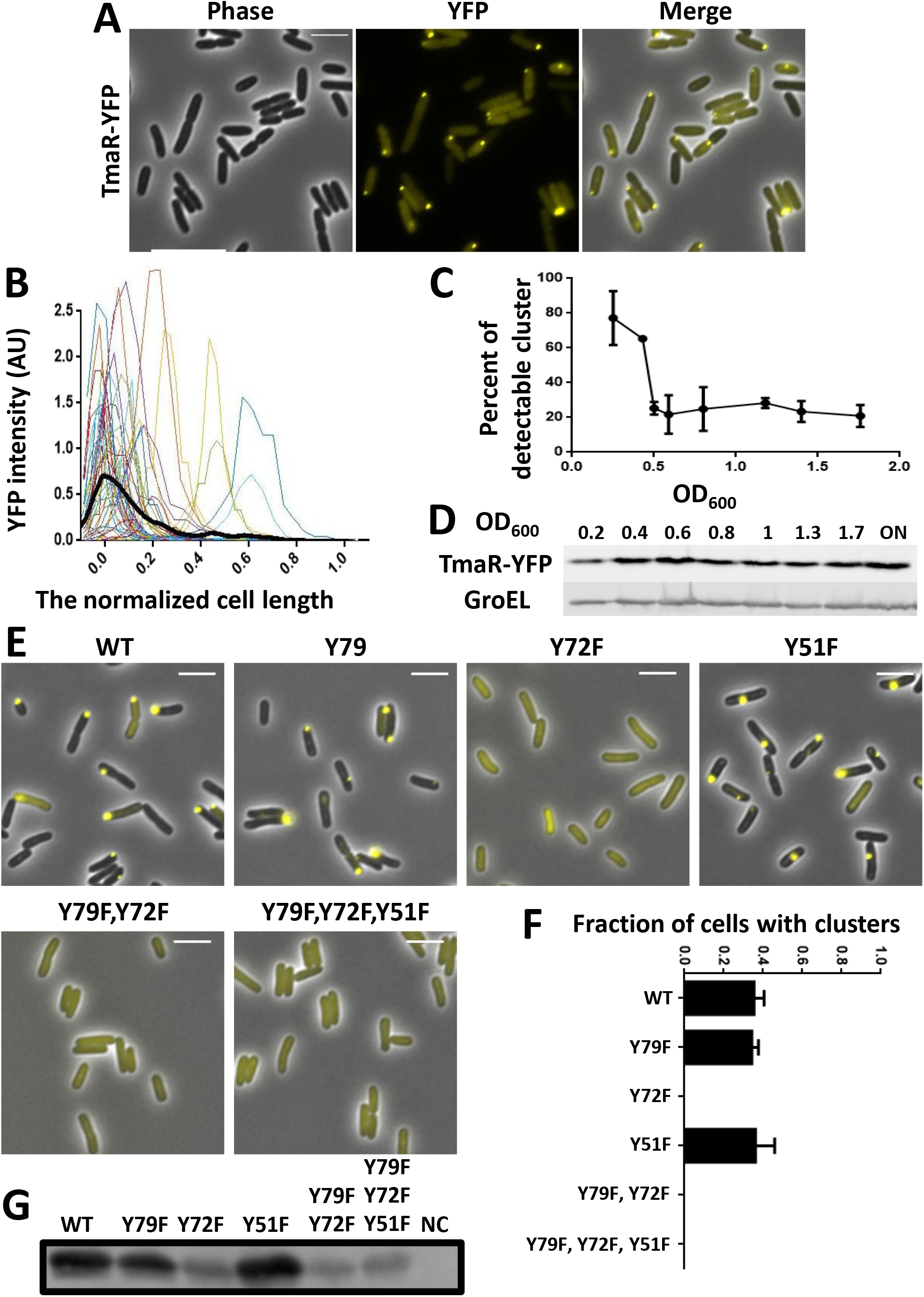
TmaR forms clusters that localize to the pole of *E. coli* cells in a tyrosine phosphorylation-dependent manner. (A) Images of cells expressing TmaR-YFP (yellow) from its native promoter and locus in the chromosome. Cells were grown in LB to OD600=0.2. Cells are in phase (gray). (B) The fluorescence intensity profile of 50 cells expressing a cluster of TmaR-YFP along the normalized long cell axis. Each cell is presented by a different color and the average is shown by a bold black line. YFP intensity is in arbitrary units. (C) Percent of cells at the indicated OD600 that showed a detectable cluster in the population. Calculation was made with the strain expressing TmaR-YFP grown in LB (N=500). The bars show the standard deviation between three biological repeats. (D) Western blot analysis showing the level of TmaR-YFP in cells at the indicated OD600 (ON, cells grown overnight) and of GroEL (control used for normalization) detected by α-mCherry and α-GroEL antibodies, respectively. (E) Images of cells expressing YFP-tagged TmaR (WT) and its variants mutated in each of its three tyrosines, (Y79F, Y72F and Y51F), in two of its tyrosines (Y79F,Y72F) and in all three (Y79F,Y72F,Y51F), all expressed from TmaR native promoter and locus in the chromosome. Cells are in phase (gray). (F) The average fraction of cells with clusters of TmaR derivatives in each strain. WT; TmaR Y79F; TmaR Y51F, TmaR Y79F, Y72F and TmaR Y79F,Y72F,Y51F. The bars show the standard deviations between different fields (N=250). (G) Western blot analysis of cells expressing YFP-tagged TmaR (WT) or its variants mutated in each of the three tyrosines, (Y79F, Y72F and Y51F), in two of the tyrosines (Y79F,Y72F) and in all three tyrosines (Y79F,Y72F,Y51F), all from the pET15b vector, as well as cells with no plasmid (NC, negative control). The membrane was probed with anti-phosphorylated tyrosine antibody.

To further characterize TmaR subcellular localization, we calculated the percentage of cells with detectable TmaR clusters in the population during growth. The results show that in LB, nearly 80% of cells had detectable TmaR clusters at their pole in early logarithmic (log) phase. This fraction dropped to 25% in mid-log phase and remained constant during stationary phase (SP) (**Fig. 1C**). To find out if the change in the fraction of cluster-harboring cells is due to a change in TmaR cellular level or to relocation of pre-existing TmaR molecules, we compared the amount of TmaR in cells at different growth stages by Western blot analysis. The results in **Fig. 1D** and its quantification in **(see SI Appendix, Fig. S3**) show that the amount of TmaR does not change much among the different growth phases. This result indicates that the growth phase-dependent difference in the percentage of cells with detectable clusters is due to a change in the distribution pattern of the TmaR molecules, i.e., whether they are assembled into clusters or diffuse in the cytoplasm, and not due to changes in TmaR cellular level.

Our studies of TmaR encouraged us to search the website for any piece of information regarding this protein. Interesting, we found that TmaR is phosphorylated on a tyrosine residue (http://dbpsp.biocuckoo.org/). This database documents the results of a high throughput analysis of the *E. coli* phosphoproteome, which identified TmaR as phosphorylated on tyrosine 79 (*21*). Because TmaR has only 3 tyrosines in positions 55, 72 and 79, we mutated each to a phenylalanine, which is the most similar amino acid to tyrosine, but cannot be phosphorylated. We thus generated strains, expressing YFP-tagged TmaR variants, Y79F, Y72F or Y51F, from their endogenous gene locus on the chromosome. Using fluorescence microscopy, we observed YFP-tagged TmaR Y79F and TmaR Y51F at the bacterial cell poles, similar to wild-type TmaR-YFP (**Fig. 1E**). When calculating the fraction of cells with a polar cluster in each of these three strains (WT and two mutants), no significant change was observed (**Fig. 1F**), indicating that the two mutants behave precisely like WT. Contrarily, the YFP-tagged TmaR Y72F mutant did not localize to the poles and was completely spread within the cytoplasm (**Fig. 1E**). Moreover no polar clusters were observed in cells expressing the YFP-tagged double mutant TmaR Y72F,Y79F or the triple mutant TmaR Y79F,Y72F,Y51F, just like in cells expressing the single mutant TmaR Y72-YFP (**Fig. 1E** and **1F**). Taken together, these results suggest that phosphorylation of TmaR on Tyr72 is a prerequisite for its polar localization.

To substantiate the above results, we monitored the localization of monomeric YFP-tagged TmaR variants and obtained similar results, i.e., wild-type TmaR and the Y79F and Y51F TmaR mutants were able to form polar clusters, while TmaR Y72F and the double and triple mutants were diffused in all cells (see SI Appendix, **Fig. S4**). To verify that the differences detected in subcellular localization is not due to a change in the cellular level of TmaR caused by the mutations, we compared the amount of mYFP-TmaR and its tyrosine mutants by Western blot analysis (similar amounts of lysates were analyzed, see SI Appendix, **Fig. S5,** lower row). The results show that the amount of TmaR does not change in the mutants (see SI Appendix, **Fig. S5**, upper panel). Finally, although there is no good mimic for phosphorylated tyrosine (pTyr), we mutated Tyr72 to glutamic acid (TmaR Y72E) or to aspartic acid (TmaR Y72D), which are both negatively charged and can partially mimic phosphorylation of other residues (*22*). The resultant mutant proteins behaved like the phenylalanine substitution, that is, they did not cluster at the pole (see SI Appendix, **Fig. S6**). This was expected, because, in terms of charge strength and geometry, glutamate and aspartate are not a good mimic for a constantly phosphorylated tyrosine (*23*). We therefore, continued to work with the Tyr to Phe substitutions, since they are supposed to have a minimal effect on the protein structure.

To get a direct proof for TmaR phosphorylation on tyrosine 72, we compared the amount of tyrosine-phosphorylated TmaR and its mutants by Western blot analysis using anti-pTyr antibody. All TmaR variants were expressed from a plasmid, because their level, when expressed from their endogenous gene locus, did not allow their unambiguous detection by this antibody and similar amounts of lysates were analyzed (see SI Appendix, **Fig. S7**). The results in **Fig. 1G** show that the three polarly localized TmaR proteins - WT, Y79F and Y51F - were nicely detected by the anti-pTyr antibody, whereas the three diffused TmaR proteins - Y72F and the double and triple mutants - showed a very faint band, frequently observed with this antibody due to phosphorylation on other residues (*24*). These results prove that Tyr72 of TmaR gets phosphorylated and, together with the imaging results, indicate that only the phosphorylated form of TmaR forms polar clusters.

Next, we aimed at identifying the kinase that controls TmaR polar localization by tyrosine phosphorylation. Two bacterial tyrosine kinases (BY kinases), Wzc and Etk, were identified in *E. coli*, the first involved in exopolysaccharide production and the second in capsule assembly (*25, 26*). To inquire whether one of these kinases phosphorylates TmaR, we transduced Δ*wzC* (Wzc-KO) or Δ*yccC* (Etk-KO) to our TmaR-YFP-expressing strain. No change in TmaR polar localization was observed as a result of either tyrosine kinase absence (see SI Appendix, **Fig. S8)**. Notably, no reduction in the cellular level of tyrosine phosphorylation was detected in a strain deleted both *wzc* and *etk* genes, indicating that *E. coli* contains yet unknown tyrosine kinases (*27*). Unless TmaR is phosphorylated by both Wzc and Etk, one of the unknown kinases is expected to phosphorylate TmaR.

### TmaR and the general PTS protein EI exhibit co-localization and co-dynamics and are capable of interacting with each other

The general PTS protein EI initiates the phosphorylation cascade that controls hierarchical uptake and utilization of preferred carbohydrates from complex environments (*28*). The soluble EI protein has previously been shown in our lab to shift between assembling into polar clusters and diffusing in the cytoplasm (*13–15*), but the factor responsible for its polar localization remained unknown (*13, 14*). We, therefore, asked whether TmaR is involved in determining the subcellular localization of EI. Imaging cells, which express TmaR-YFP and EI-mCherry expressed from their respective native promoters and loci on the chromosome, by fluorescence microscopy demonstrated that the polar clusters of the two proteins co-localize (**Fig. 2A**).

**Figure 2.**
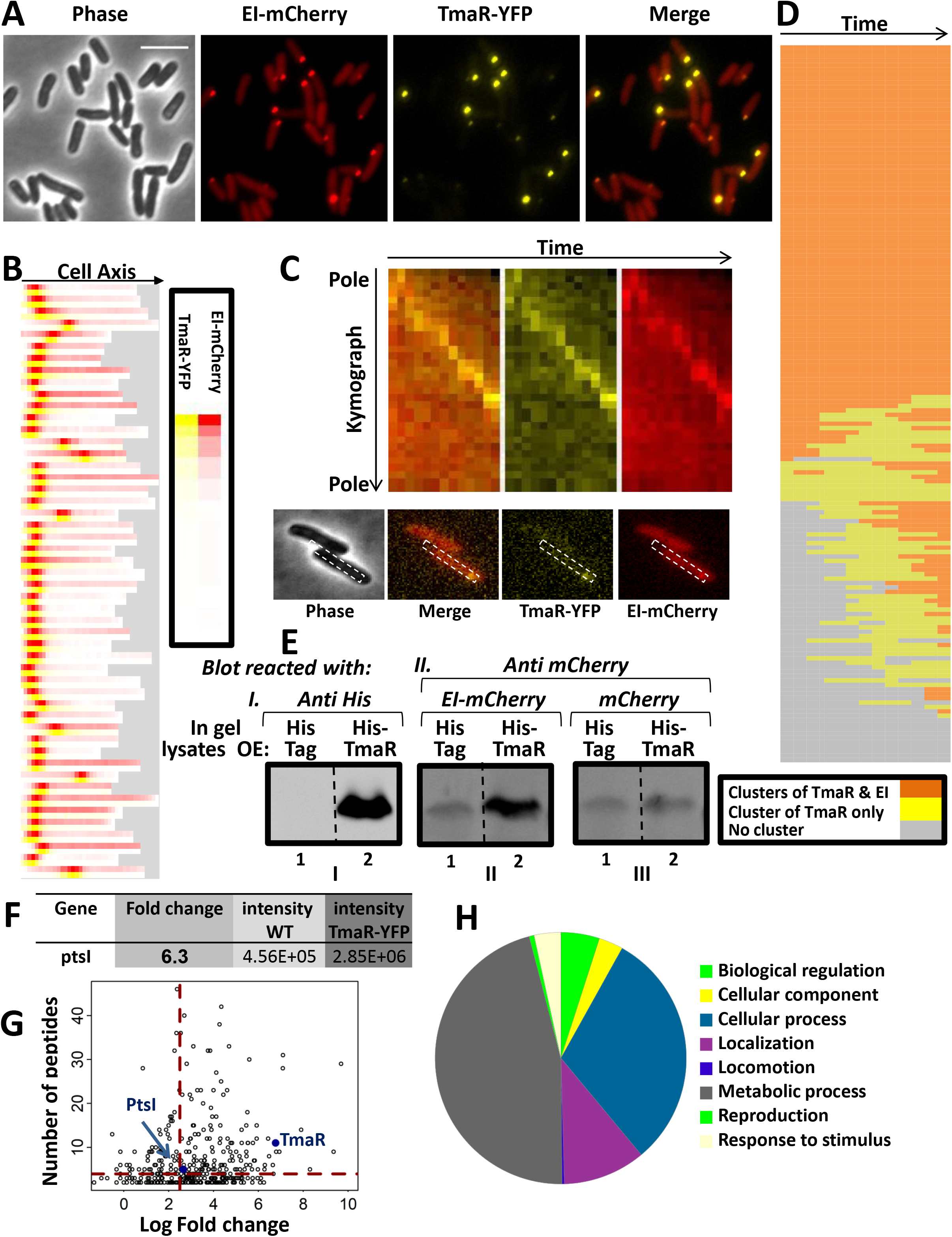
TmaR and EI exhibit co-localization and co-dynamics and are capable of interacting with each other. (A) Images of cells expressing EI-mCherry (red) and TmaR-YFP (yellow), both expressed from their native promoter and locus in the chromosome. Cells are in phase (gray). (B) Heat map showing the localization of EI-mCherry (red) and TmaR-YFP (yellow). The color intensity represents the strength of the fluorescent signal in the same cell along the long cell axis (arbitrary units, AU). Each lane represents a single cell (N=100). Because of the heterogeneity in cells length, the area that is devoid of cell body is in grey. (C) Upper panels: Single-cell kymograph of the merged TmaR-YFP and EI-mCherry (orange), TmaR-YFP (yellow) and EI-mCherry (red). The kymograph X-axis is time (0-12 min) and the Y-axis is the cell length along its long axis. Lower panels: The cell (outlined) shown in the kymograph. (D) TmaR-YFP and EI-mCherry clusters were monitored in 163 cells every 3 minutes for 3 hours by time-lapse microscopy. Each row represents one cell over time. Orange, both TmaR and EI clusters are present in the cell; yellow, only the TmaR cluster is present in the cell; grey, no cluster is present in the cell. (E) Far-Western analysis of the interaction between EI and TmaR. Lysates of Δ*tmaR* cells expressing the His-tag only (lanes 1) or His-tagged TmaR (lanes 2) were analyzed by SDS-PAGE, blotted onto a nitrocellulose membrane and probed with: anti-His antibody (**I**), EI-mCherry followed by anti-mCherry antibody (**II**) or mCherry followed by anti-mCherry antibody (**III**). The observed bands match His-TmaR size (~15 kDa). (F) Mass spectrometry analysis showing the fold change of EI co-purification on anti-GFP antibody-bound resin from cells endogenously expressing TmaR-YFP vs. cells endogenously expressing non-tagged TmaR (WT). The *ptsI* gene codes for the EI protein. (G) Volcano plot presentation of the mass spectrometry results. The number of peptides identified for each protein was plotted against the log fold change in this protein intensity between cells expressing TmaR-YFP and cells expressing non-tagged TmaR (WT). The EI (PtsI) and TmaR are depicted as blue dots. A red dotted lines show the values for EI. (H) A pie chart showing the biological processes, identified by GO analysis, in which the proteins identified as significantly co-purifying with TmaR by mass spectrometry are involved.

Although EI-mCherry activity has been demonstrated previously (*13*) and shown to be comparable to that of wild-type EI (*15*), not only did we verify this again (see SI Appendix, **Fig. S9**), but we also asked if EI-mCherry fluorescent signal correlates with EI-mCherry level. To this end, we expressed EI-mCherry from a plasmid by increasing concentrations of inducer and measured the mean intensity of the EI-mCherry fluorescent signal by imaging (see SI Appendix, **Fig. S10A**), as well as the protein level using Western blot analysis (see SI Appendix, **Fig. S10B,C**). These results show that the fluorescence intensity of EI-mCherry is proportional to the level of the EI protein. Of note, a higher fraction of EI-mCherry diffuse molecules was detected when it was co-expressed with non-tagged TmaR compared to when it was co-expressed with TmaR-YFP (see SI Appendix, **Fig. S11A**). However, the fraction of cells with a polar EI-mCherry cluster in populations expressing YFP-tagged TmaR is only slightly lower than when expressed with non-tagged TmaR (see SI Appendix, **Fig. S11B**). Although these differences are minor and are not expected to affect our conclusions, we refrained from observing EI localization in the presence of fluorescently-tagged TmaR, unless we aimed at exploring their colocalization.

To assess the degree of EI and TmaR co-localization, we inspected the localization of EI-mCherry and TmaR-YFP along the long cell axis in 100 cells that contained clusters of both proteins, which represented 63% of the population. The results, presented as a heat map in **Fig. 2B**, show that the maximal fluorescence of the two proteins overlaps in all cells and that the two proteins mostly localize to one pole, indicating co-localization of TmaR and EI clusters. Because the clusters of both proteins exhibited dynamics, mainly within the pole region, we asked whether they move together or independently. The spatiotemporal behavior of co-expressed YFP-TmaR and EI-mCherry over time, recorded by time-lapse microscopy, indicated that the dynamics of the two proteins is coordinated, as shown by the kymograph in **Fig. 2C**. To study the co-occurrence of TmaR and EI clusters, we spotted diluted overnight culture of cells expressing YFP-TmaR and EI-mCherry on an agar pad with fresh medium and monitored the existence of TmaR and EI clusters in 163 cells every 3 minutes for 3 hours by time-lapse microscopy. The vast majority of the cells divided once during this period (growth in agar chambers of minimal media without aeration is rather slow) and images of both daughter cells were analyzed, culminating in a final population of about 320 cells that were finally analyzed. The results, presented in **Fig. 2D**, can be summarized as following: 79% of the cells contained both TmaR and EI clusters, approximately two thirds contained clusters of the two proteins throughout their lifetime, and the rest contained clusters of both at a certain point during their life. Notably, none of the cells in the population contained an EI cluster without a TmaR cluster at any point, whereas 15% of the cells contained only a TmaR cluster sometime during their lifetime. The remaining 6% did not contain any cluster at any time. These results imply that TmaR clusters in the vast majority of *E. coli* cells, throughout their lifetime or temporarily, with a significant fraction of the cells having also an accompanying EI cluster.

Co-localization and co-dynamics of TmaR and EI raised the possibility that the two proteins interact with each other. To check this possibility, we asked if TmaR and EI can be co-purified from cells. To this end, we lysed cells expressing or not endogenously expressed TmaR-YFP and incubated the lysates with anti-GFP antibody-bound beads. The proteins that bound to the beads from each strain were eluted and subjected to mass spectrometry analysis, and the fold change of YFP-TmaR binding intensity compared to the non-tagged TmaR intensity was calculated for each protein. EI was detected as co-purifying with TmaR at a high (6.3) fold change (**Fig. 2F**). When plotting the number of peptides versus the log fold change for all the proteins detected in the mass spec analysis (**Fig. 2G**), EI comes up as a bona fide interactor. A Gene Ontology (GO) analysis of the mass spec results identified the biological processes that TmaR-interactors are involved in (**Fig. 2H**). Notably, the biggest group of TmaR partners is of proteins involved in metabolic processes, raising the possibility that by localizing EI, TmaR is involves in its regulation. Of note, another big group of TmaR partners is of proteins that are associated with localization.

Because co-purification of TmaR and EI might have been due to indirect interaction, we used the Far-Western technique to ask whether the two proteins can interact directly. To this end, lysates of Δ*tmaR* cells expressing His-tagged TmaR from a plasmid (**Fig. 2E**, lanes 2) and a control lysate of Δ*tmaR* cells expressing only the His tag (**Fig. 2E**, lanes 1) were subjected to SDS–PAGE and blotted onto a nitrocellulose membrane. Three identical membranes (I-III) with identical amount of the two lysates were prepared and probed with anti-His antibody (Western, **Fig. 2E**, panel I) or with purified EI-mCherry or mCherry, followed by antibodies against mCherry (Far-Western, **Fig. 2E** and **see SI Appendix, Fig. S12**, panels II and III). The Far-Western results show that EI-mCherry, but not mCherry, interacted with TmaR-His, indicating that TmaR and EI have the capacity to interact directly. Of note, the faint band that co-migrates with TmaR, observed in all lanes in membranes II and III, is a non-specific band, as it appears regardless of the lysate analyzed, and it could be explained by the use of polyclonal antibody or by non-specific interaction with mCherry.

Together the results in this section suggest that TmaR and EI co-assemble at the cell pole.

### TmaR controls EI polar localization, but not *vice versa*

The results in the previous section indicate that the subcellular distribution of TmaR and EI is coordinated. To test whether one of the two proteins affects localization of the other or their localization is co-dependent, we first observed the localization of EI-mCherry in wild-type (WT) and Δ*tmaR* (TmaR-KO) strains, as well as in a strain overexpressing TmaR (TmaR-OE). Deletion of the *tmaR* gene had a dramatic effect on EI localization pattern, that is, the cells contained no detectable clusters and EI-mCherry was observed as homogenously diffused in the cytoplasm (**Fig. 3A** and **3B**, compare WT and TmaR-KO), indicating that EI polar localization depends on TmaR. To substantiate that TmaR is the cause for polar localization of EI clusters, we complemented the TmaR-KO strain by expressing TmaR from a plasmid and monitored EI-mCherry localization. The results demonstrated that polar localization of EI clusters was restored (**Fig. 3A**, TmaR-KO+OE). Of note, the population of wild-type cells overexpressing TmaR contained twice the number of cells with EI-mCherry polar clusters in comparison to the population of cells expressing TmaR from the chromosome (**Fig. 3B**, compare WT and TmaR-OE). Moreover, a comparison of the sum intensity of the EI-mCherry clusters fluorescent signals suggested that the average EI-mCherry cluster in the TmaR-OE strain contains significantly more EI-mCherry molecules than the strain expressing TmaR from the chromosome (**Fig. 3C**). These results imply that the ability of EI to localize to the pole depends entirely on TmaR and the extent of EI recruitment to the pole correlates with the cellular level of TmaR.

**Figure 3.**
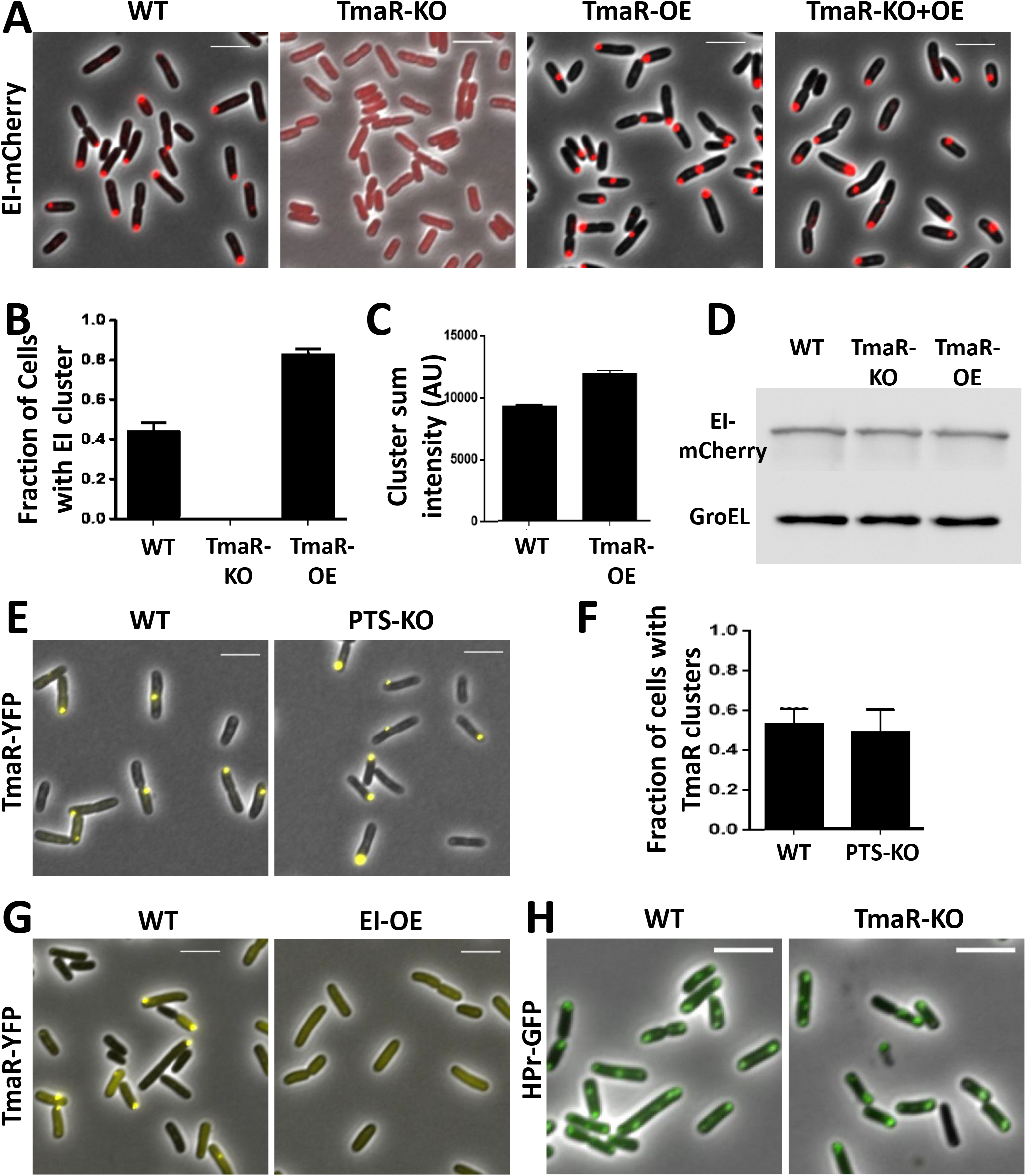
TmaR is essential for EI polar localization, but not *vice versa*. (A) From left to right: Images showing EI-mCherry in wild-type cells (WT), in cells deleted for the *tmaR* gene (TmaR-KO), in wild-type cells overexpressing TmaR (TmaR-OE) and in TrmaR-KO cells overexpressing TmaR (TmaR-KO+OE). (B) The average fraction of cells with EI clusters in the different strains: WT, TmaR-KO and TmaR-OE (N=500). The bars show the standard deviation between different fields. (C) The mean of the sum intensity of the EI-mCherry clusters (arbitrary units, AU): WT and TmaR-OE (N=464). The bars show the standard deviation between different fields. (D) Western blot analysis showing the level of EI-mCherry and GroEL proteins (control used for normalization) detected by α-mCherry and α-GroEL antibodies, respectively. (E) Images showing TmaR-YFP localization in wild-type cells (WT, left panel) and in cells deleted for the *pts* operon that encodes EI (PTS-KO, right panel). TmaR-YFP was expressed from its native promoter and locus in the chromosome. Cells are in phase (grey). (F) The average fraction of cells with TmaR clusters in WT and PTS-KO strain (N=400). The bars show the standard deviations between different fields. (G) Images showing the localization of TmaR-YFP, expressed from the chromosome, in wild-type cells (WT, left panel) and in cells overexpressing EI (EI-OE). (H) Images of wild-type cells (WT, left panel) and cells deleted for *tmaR* (TmaR-KO, right panel) overexpressing HPr-GFP. Cells are in phase (grey).

The observed differences in EI-mCherry localization in the WT, TmaR-KO and TmaR-OE strains is not due to differences in EI cellular level, as indicated by the Western blot analysis results presented in **Fig. 3D**, which show that the three strains contain comparable amounts of EI-mCherry. Hence, pre-existing EI molecules can join the cluster or diffuse out of it.

Next, we asked whether polar localization of TmaR depends on EI localization by observing TmaR-YFP expressed from the chromosome of cells deleted for the *pts* operon that encodes EI (PTS-KO). The results in **Fig. 3E** show that TmaR was not affected by the absence of EI, as localized to the pole of these cells. Furthermore, we did not observe a significant change in the fraction of cells with TmaR clusters between the PTS-KO and the wild-type strains (**Fig. 3F**). Hence, polar localization of EI requires TmaR, but the opposite is not true. Still, in cells overexpressing EI from a plasmid, the chromosomally-encoded TmaR became more diffused (**Fig. 3G**). The effect of expressing EI at an artificially high concentration on TmaR distribution, which can be described as *in vivo* titration, supports the notion of physical interaction between the two proteins.

EI is the first general PTS protein, which initiates the cascade that controls carbohydrate utilization by phosphorylating the second general PTS protein, HPr. Although both EI and HPr exhibit polar localization, they were shown to localize to the poles independently (*13*). We, therefore, asked whether TmaR also controls HPr polar localization. The results in **Fig. 3H** show that polar localization of HPr-GFP was not affected when expressed in a TmaR-KO background, indicating that TmaR specifically controls EI, but not HPr localization.

### TmaR-mediated EI localization affects sugar uptake by the PTS system

The results presented above show that TmaR has a crucial role in EI polar clusters localization, raising the possibility that it affects EI function in PTS sugars uptake. To test this proposition, we first streaked TmaR-WT, TmaR-KO and TmaR-OE strains on MacConkey plates supplemented with different PTS sugars, with a PTS-KO strain serving as a negative control. **Fig 4A** shows growth of these strains on a fructose-supplemented plate, but similar results were observed with MacConkey plates supplemented with glucose, mannitol or N-acetylglucosamine (NAG). In all cases was the red color of the colonies of the TmaR-KO strain more intense than that of the WT colonies, and the red color of the WT deeper than that of the TmaR-OE strain. These results suggest that the absence of TmaR makes cells hyper-active for PTS sugar consumption, while overexpression of TmaR reduces PTS activity.

**Figure 4.**
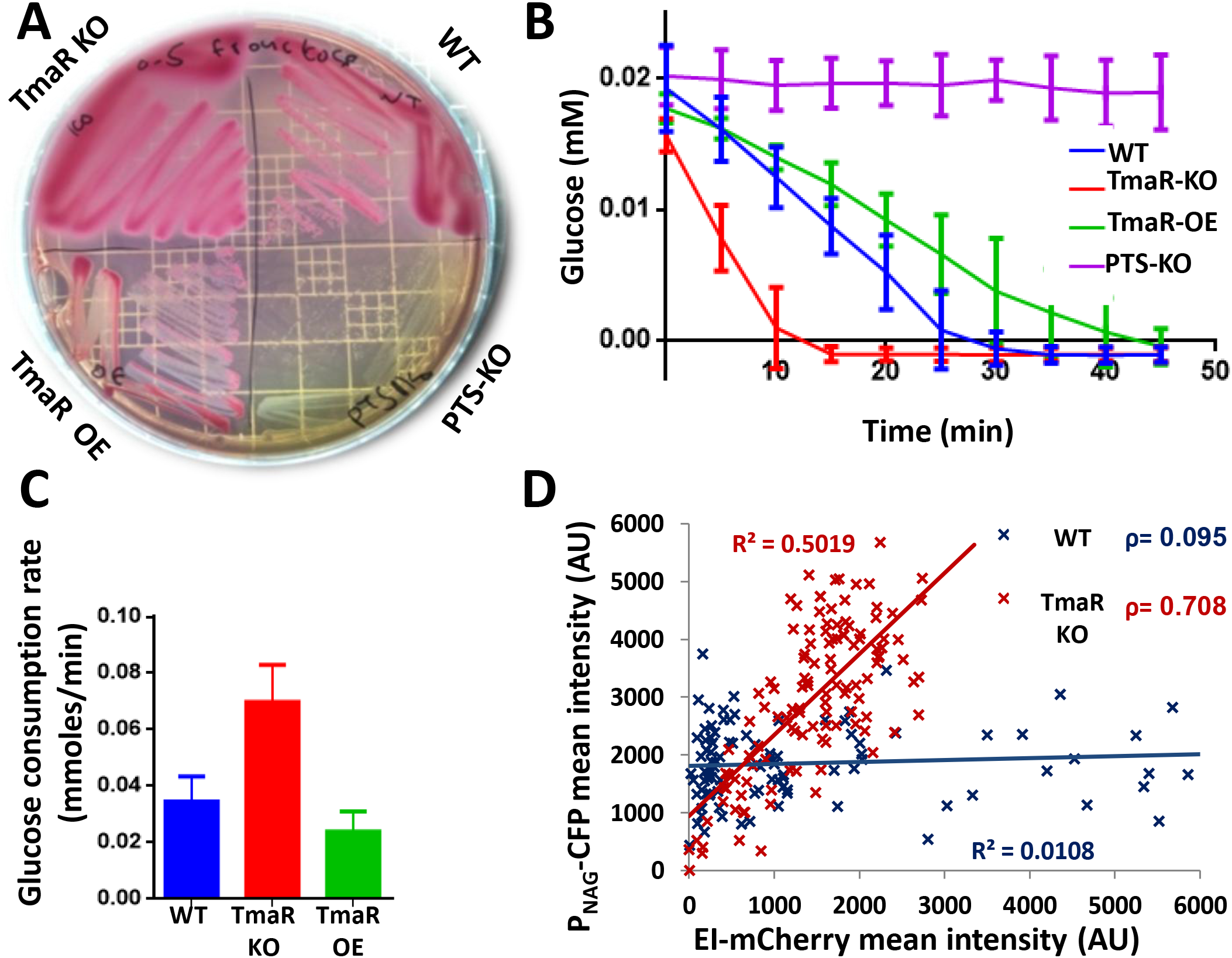
TmaR-mediated EI localization affects sugar uptake by the PTS system. (A) A representative MacConkey plate supplemented with 0.4% fructose showing the phenotype of wild-type cells (WT), cells deleted for the *tmaR* gene (TmaR-KO), wild-type cells overexpressing TmaR (TmaR-OE) and cells deleted for the *pts* operon (PTS-KO). (B) The rate of glucose consumption by the following strains: WT (blue), TmaR-KO (red) TmaR-OE (green), and PTS-KO (purple). The bars show the standard deviation between three biological repeats. (C) The rate of glucose consumption in mmoles per min for the WT (blue), TmaR-KO (red) and TmaR-OE (green). The bars show the standard deviation between three biological repeats. (D) A scatter plot showing the mean intensity (arbitrary units, AU) of mCerulean expressed from *P_NAG_* as a function of the cellular EI-mCherry mean intensity (AU) after 1 hour of transition to NAG-continuing medium for WT cells (blue Xs, N=103) and TmaR-KO strain (red Xs, N=103). The r squared (R^2^) values and Pearson correlation coefficients (ρ) are given.

Because TmaR-YFP was used for the TmaR-EI localization and interaction studies, we verified that fluorescently tagging TmaR does not affect EI function by streaking cells expressing TmaR-YFP, mYFP-TmaR or untagged TmaR on a fructose-supplemented MacConkey plate. The results show that EI is similarly active in the three strains (see SI Appendix, **Fig. S13**).

To quantify the effect of TmaR on EI function, we compared the ability of strains expressing TmaR-WT, TmaR-KO and TmaR-OE to consume glucose by monitoring the drop in glucose concentration in the medium over time (see Supporting Materials and Methods). The results in **Fig. 4B** verified that the TmaR-KO strain consumes glucose faster than the WT strain, which consumes glucose faster than the TmaR-OE strain. Comparison of the average consumption rate of the three strains corroborated that TmaR-KO consumes glucose two times faster than the WT and TmaR-OE strains (**Fig. 4C**). Hence, the activity of EI inversely correlates with the level of TmaR in the cell. These results strengthen our previous results, which suggested that clustering of EI negatively correlates with EI activity (*15*).

The increased sugar consumption by the Δ*tmaR* mutant could be caused by increased availability of EI or it could be caused by increased levels of the glucose transporter protein PtsG (EIICB^Glc^), shown before to affect glucose consumption (*29*). To discriminate between these possibilities, we compared the amount of PtsG in WT and TmaR-KO cells by Western blot analysis, using antibodies against the cytoplasmic C-terminal domain of PtsG (anti-IIB^Glc^ antibodies, (*30*)). We saw no difference in the level of PtsG between the two strains, which was lower than PtsG level in cells overexpressing it from a plasmid (see SI Appendix, **Fig. S14**). These results support the notion that the increase in sugar consumption in cells lacking TmaR is due to an increase in EI activity.

An additional assay that we used to compare the effect of TmaR subcellular level on PTS sugars uptake relied on the use of a strain expressing mCerulean, a cyan fluorescent protein (CFP), from the N-acetyl glucosamine (NAG) promoter (P_*NAG*_) and EI-mCherry (*31*). NAG has been used in the past as a representative of PTS sugars, whose utilization is positively regulated by EI, since induction of *P_NAG_* by NAG is consistent with the positive feedback mechanism suggested for promoters of PTS sugar utilization genes (*31*)(*15*). Hence, production of mCerulean from *P_NAG_* implies that EI is active and enables estimation of EI activity by fluorescence microscopy. After transducing the TmaR-KO mutation into the EI-mCherry and *Pnag*-mCerulean expressing strain, the single cell fluorescence mean intensity of mCerulean and of EI-mCherry, which report on EI activity (see SI Appendix, **Fig. S10**) and on EI cellular level, respectively, were plotted against each other for 103 TmaR-WT cells and 103 TmaR KO cells grown in minimal medium containing NAG. The results in **Fig. 4D** show lack of correlation between the amount of EI and its activity in the TmaR-WT cells, whereas the level of EI in the TmaR-KO strain correlated with its function, as indicated by the Pearson correlation scores (p) and the r squares (R^2^). These results suggest that in the strain expressing wild-type TmaR, the amount of active EI does not correlate with its cellular level, but rather with the fraction that is not sequestered at the pole. Conversely, in the TmaR-KO strain, in which all EI molecules are diffuse, the activity of EI correlates with the level of EI molecules in the cell. These results are in agreement with the sugar consumption assay (**Fig. 4B**), which showed that EI activity inversely correlates with TmaR level. When presenting the fluorescence mean intensity of mCerulean by a box plot, the level of this protein production in TmaR-KO cells emerges as more heterogeneous than in WT cells (see SI Appendix, **Fig. S15)**, suggesting that not only does TmaR control EI activity, but it also limits the degree of heterogeneity in its activity, a phenomenon that has been studied in our lab (*15*). This implies that flooding the cells with excess of EI, which allows irregular level of sugar consumption, disturbs the cells sense of balance and leads to higher fluctuations in sugar uptake.

Overall, the results in this section demonstrate that there is a connection between EI localization and function. That is, in the absence of TmaR, EI is diffuse and, therefore, more active in sugar uptake, whereas upon TmaR expression, a fraction of EI assembles into polar clusters and, therefore, less sugar is taken by the cells. Moreover, our results hint at the possibility that TmaR-mediated EI assembly controls the population heterogeneity with respect to sugar consumption.

### Tyrosine phosphorylation of TmaR is required for the release of EI from the poles and for its activity

The results described thus far, which show that TmaR controls EI localization and function and has the potential to interact with EI, made us wonder whether TmaR phosphorylation on Tyr72, which is required for its polar localization, affects its ability to interact with EI and to control EI localization and function.

We first tested if EI interaction with TmaR depends on TmaR tyrosine phosphorylation using the Far-Western technique. To this end, similar amounts of lysates ((see SI Appendix, **Fig. S16A)** of Δ*tmaR* cells expressing wild-type TmaR, the single mutants in TmaR tyrosines (Y79F and Y51F), the double mutant or the triple mutant (see above), all from plasmids as His-tagged proteins, as well as lysates of Δ*tmaR* cells expressing only a His-tag or no protein from plasmids as controls, were subjected to SDS–PAGE. The proteins were blotted onto a nitrocellulose membrane, incubated with purified mCherry-tagged EI, and then with antibodies against the mCherry. The Far-Western results (**Fig. 5Ab**), before and certainly after quantification by normalizing them to the Western results (**Fig. 5Aa** and **5c**), clearly show that although EI can interact with all TmaR variants, the interaction with TmaR proteins that lack Tyr72 (non-phosphorylated) is significantly stronger. These results indicate that EI affinity for unphosphorylated TmaR is much higher than its affinity for phosphorylated TmaR.

**Figure 5.**
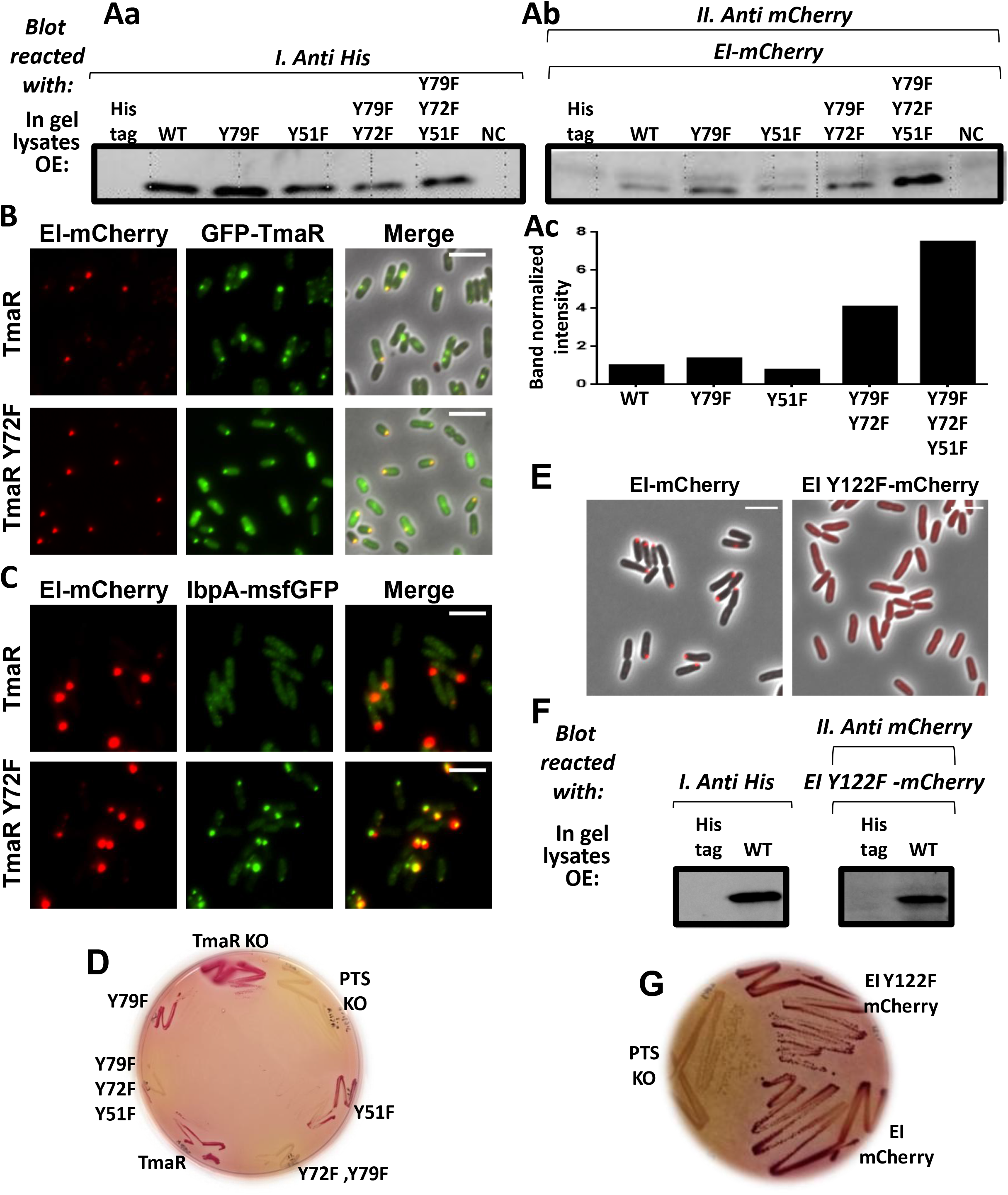
The effect of replacement of TmaR and EI tyrosines on their interaction and on EI localization and activity. (A) Far-Western analysis of the interaction between purified EI and TmaR (WT) or its variants mutated in each of the three tyrosines, (Y79F, Y72F and Y51F), in two of the tyrosines (Y79F,Y72F) and in all three tyrosines (Y79F,Y72F,Y51F), all expressed from plasmids as His-tagged proteins in a Δ*tmaR* background, as well as lysates of Δ*tmaR* cells expressing only a His-tag or no protein from plasmids as controls (NC, negative control). Equal amount of lysates were fractionated by SDS-PAGE and blotted onto two nitrocellulose membranes, one incubated with anti-His antibodies (Western, **a**) and the other with purified mCherry-tagged EI followed by antibodies against mCherry (Far Western, **b**). The detected band is at TmaR expected size (~15 kDa). The intensity of the bands detected in the Far Western analysis was normalized to the intensity of the respective bands detected in the Western blot analysis (**c**). (B) Images of cells showing localization of EI-mCherry in cells overexpressing wild-type GFP-TmaR (TmaR) or non-phosphorylated GFP-TmaR (TmaR Y72F). (C) Images of cells expressing EI-mCherry and IbpA-msfGFP in cells overexpressing non-tagged wild-type TmaR (TmaR) or non-phosphorylated TmaR (TmaR Y72F). (D) A representative MacConkey plate supplemented with 0.4% fructose and 0.1 mM IPTG showing the phenotype of Δ*tmaR* cells overexpressing TmaR (WT) and its mutants with the indicated substitutions of TmaR tyrosines, all expressed from the *lac* promoter in pCA24N, as well as of Δ*tmaR* (TmaR-KO) cells and cells deleted for the *pts* operon (PTS-KO).. (E) Images of cells expressing EI-mCherry and EI Y122F-mCherry, both expressed from the *ptsI* promoter and locus in the chromosome. Cells are in phase (gray). (F) Far-Western analysis of the interaction between purified EI Y122F-mCherry and His-tagged TmaR in a Δ*tmaR* background. The details are as described in Fig. 5A for wild-type EI. The proteins were blotted onto two nitrocellulose membranes, one incubated with anti-His antibodies (Western, left panel) and the other with purified mCherry-tagged EI Y122F followed by antibodies against mCherry (Far Western, right panel). (G) A representative MacConkey plate supplemented with 0.4% fructose showing the phenotype of cells expressing EI-mCherry and EI Y122F-mCherry from *ptsI* promoter and locus on the chromosome and of cells deleted for the *pts* operon (PTS-KO).

The increased binding of non-phosphorylated TmaR to EI raised the question whether TmaR phosphorylation on Tyr72, which prevents its polar clustering, affects EI localization. Using fluorescence microscopy, we monitored EI-mCherry localization in the presence of plasmid-expressed wild-type GFP-TmaR or GFP-TmaR Y72F. Surprisingly, EI was recruited to the poles in both strains (**Fig. 5B**, left panels). Moreover, a portion of the GFP-TmaR Y72F mutant, clustered (**Fig. 5B**, middle panels) and co-localized with EI-mCherry (**Fig. 5B**, right panels), raising the possibility that the strong binding of EI to non-phosphorylated TmaR led to their co-aggregation and the formation of inclusion bodies. To test this hypothesis, we expressed IbpA, a reporter for inclusion bodies presence (*32*), tagged with msfGFP, together with EI-mCherry, both from the chromosome, and non-tagged TmaR or TmaR Y72F, both from a plasmid. The results show that IbpA--msfGFP formed polar foci that overlapped with EI-mCherry in cells expressing non-phosphorylated TmaR, as opposed to IbpA-msfGFP diffuse distribution in the presence of wild-type TmaR (**Fig. 5C**), indicating that EI aggregates to form inclusion bodies only in the presence of non-phosphorylate TmaR. The quantitative significance of this result is shown in SI Appendix, **Fig. S17**.

Taken together, the results thus far indicate that EI binding to TmaR is reversible as long as TmaR can get phosphorylated, whereas its binding to non-phosphorylated TmaR is very strong and quite irreversible. Hence, only phosphorylated TmaR, which recruits EI to the poles, can release EI from the polar clusters.

Next, we asked if the function of EI in sugar uptake is affected by its inability to be released from the poles when TmaR cannot get phosphorylated. To test this, we streaked Δ*tmaR* strains harboring plasmids expressing wild-type TmaR, the three single substitution of TmaR tyrosines, and the triple replacement of all TmaR tyrosines, as well as the Δ*tmaR* and Δ*pts* controls, on a fructose-supplemented MacConkey plate. The results in **Fig. 5D** show that only TmaR proteins that cannot get phosphorylated (the double and the triple mutant) are defective in sugar utilization (white colonies), whereas TmaR proteins that can get phosphorylated (WT, Y79F and Y51F) can utilize the sugar (red colonies). These results indicate that overexpression of non-phosphorylated TmaR proteins (lack Y72 residue), which results in irreversible binding of EI and aggregation, prevents EI-mediated sugar consumption. Of note, when strains expressing the above TmaR proteins from *tmaR* native promoter and locus on the chromosome where streaked on a similar plate, all TmaR mutant strains demonstrated a certain ability to consume sugar, although less than TmaR WT strain (see SI Appendix, **Fig. S18**). Hence, native expression level of nonphosphorylated TmaR leave a certain fraction of EI molecules unbound and, hence, active; however, when overexpressed, nonphosphorylated TmaR binds an increased fraction of EI and aggregates with it, thus preventing sugar consumption.

### Tyrosine phosphorylation of EI is important for its polar localization, but not for its function

Besides its well-known self-phosphorylation on His 189, a high throughput analysis (http://dbpsp.biocuckoo.org/) identified EI as a protein that undergoes phosphorylation on a tyrosine residue, Tyr122 (*27*), which is a conserved residue (*33*). To examine if Tyr122 is important for EI polar localization, we mutated this tyrosine to phenylalanine and compared the cellular distribution of EI Y122F to that of wild-type EI, after tagging both proteins by mCherry. The results in **Fig. 5E** show that unlike EI-mCherry, which was detected in polar clusters, EI Y122F-mCherry was completely diffused throughout the cytoplasm. Notably, EI Y122F does not affect TmaR localization (see SI Appendix, **Fig. S19**), as expected from the results reported above, which demonstrated that EI is not required for TmaR polar localization. The two known tyrosine kinases of *E. coli* do not seem responsible for EI phosphorylation, since transduction of a strain expressing EI-mCherry with deletions of their genes (Wzc-KO or Etk-KO) did not affect EI localization (see SI Appendix, **Fig. S20**). Hence, the phosphorylated tyrosine in position 122 of EI is important for its polar localization.

Since TmaR is important for EI recruitment to the poles, we asked whether EI Y122F non-polar localization is due to its inability to interact with TmaR. Using the Far-Western technique, we tested the ability of EI Y122F to interact with TmaR. For this purpose, similar amounts of lysates of Δ*tmaR* cells expressing His-tagged TmaR or only a His-tag, were subjected to SDS–PAGE. The proteins were blotted onto two nitrocellulose membranes that were incubated either with anti-His antibodies (Western, **Fig. 5F,** left panel) or with purified EI Y122F-mCherry followed by anti-mCherry antibodies (Far Western, **Fig. 5F**, right panel). The results show that the Y122F mutation in EI did not eliminate the protein capacity to interact with TmaR.

In light of the effect of the Y122F mutation in EI on its localization, and the lack of effect of the same mutation on its ability to interact with TmaR, we asked whether this mutation affects EI function, by streaking EI-mCherry and EI Y122F-mCherry on fructose-supplemented MacConkey plates together with a PTS-KO strain (negative control). The results in **Fig. 5G** show that sugar consumption is not affected by the Y122F mutation in EI, suggesting that TmaR phosphorylation on tyrosine is not important for the function of EI in sugar uptake.

Overall our results show that tyrosine phosphorylation of both TmaR and EI is important for their polar localization. However, whereas TmaR tyrosine phosphorylation site is important for its interaction with EI, the tyrosine phosphorylation site on EI is not important for its interaction with TmaR and, therefore, does not affect its activity. Together, our results suggest that the interaction between the TmaR and EI is the crucial stage in the control EI activity as a sugar regulator.

### TmaR is important for *E. coli* survival in mild acidic conditions in a sugar- and EI-dependent manner

Our finding that a strain deleted for *tmaR* consumes glucose faster than a strain carrying the *tmaR* gene calls the role of the TmaR protein into question: under which environmental conditions that *E. coli* encounters does TmaR absence confer a disadvantage?

In nature, *E. coli* often encounters mild acidic conditions (pH~5.0), e.g., in macrophages, in the intestine and in specific foods, or extreme acidic conditions, like in mammalian stomach (pH 1-3) (*34*) (*35*). We, therefore, compared the survival of wild-type and TmaR-KO cells grown at different pH in minimal media supplemented with glucose. When grown at pH 5.5 in the presence of the most frequently used glucose concentration (0.4%), TmaR-KO displayed a drastic reduction in survival compared to the wild-type strain, as evident by the drop in the number of TmaR-KO cells after 3 days in this medium by four orders of magnitude compared to wild-type (**Fig. 6A**). The same effect was observed when cells were grown at pH 5 (**Fig. 6B**, middle panel). However, at pH 7.0 or pH 4 and below, both strains survived similarly or did not survive at all, respectively (**Fig. 6B,** upper and lower panels, respectively), indicating that only mild acidity causes the observed drop in survival of cells lacking TmaR. The reduction in survival of TmaR-KO cells in acidic medium could be reversed by inoculating the cells into a neutral pH medium after 3 days, but was irreversible after 4 days (see SI Appendix, **Fig. S21)**.

**Figure 6.**
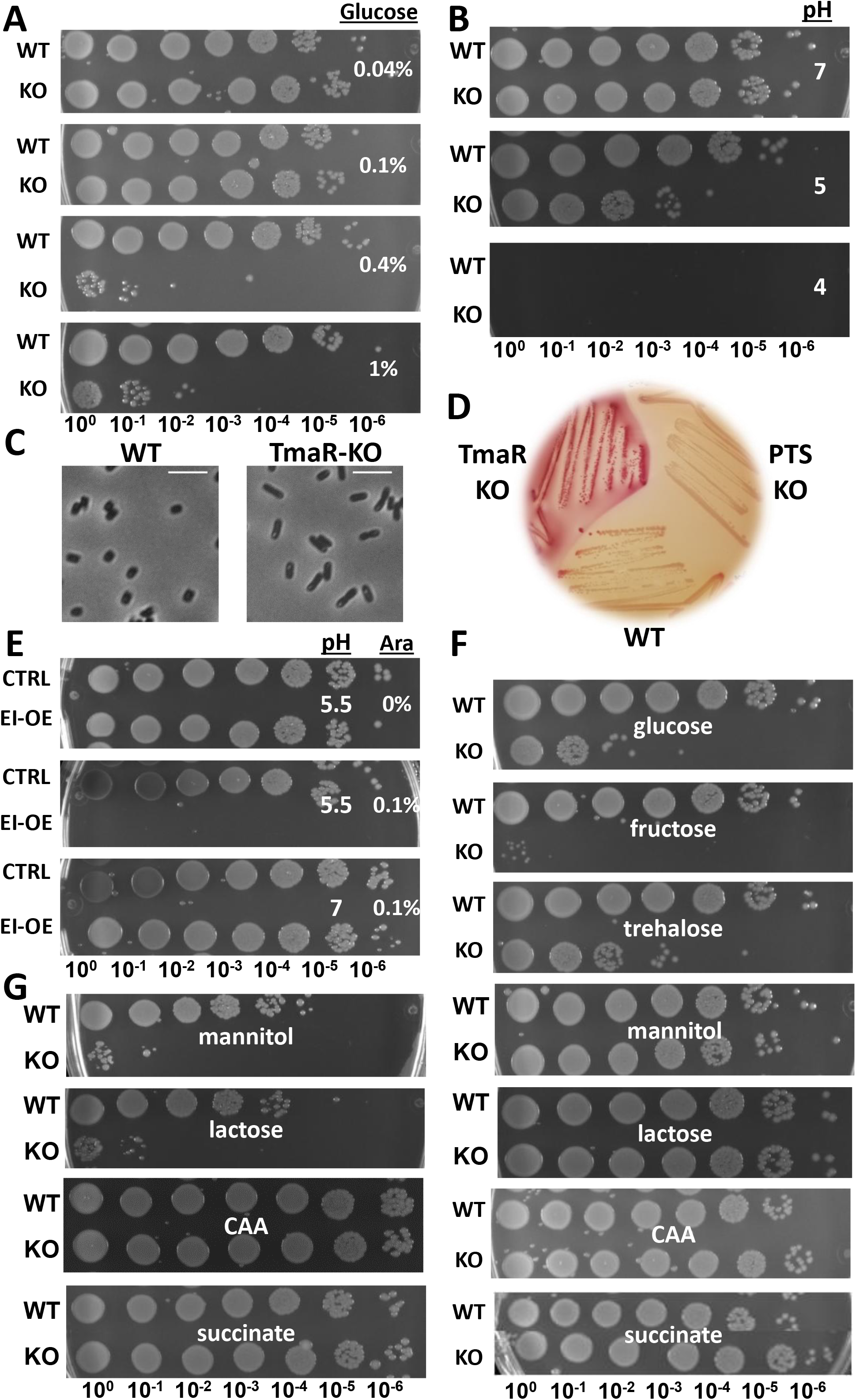
Cells lacking TmaR exhibit a drastic reduction in survival in mildly acidic conditions. (A) Pictures of wild-type (WT) and TmaR-KO (KO) colonies grown in acidic M9 medium (pH 5.5) supplemented with 0.04%, 0.1%, 0.4% and 1% glucose for 72 hours, spotted after serial dilutions on LB plates. (B) Pictures of wild-type (WT) and TmaR-KO (KO) colonies grown in acidic media adjusted to pH 7 or in neutral M9 adjusted to pH 5 or 4, all supplemented with 0.4% glucose, grown for 72 hours and spotted after serial dilutions. (C) Images of WT and TmaR-KO cells that grew in acidic M9 medium (pH 5.5) supplemented with 0.4% glucose for 72 hours. Cells are in phase. (D) A representative MacConkey plate showing the phenotype of cells expressing wild-type TmaR (WT) or TmaR-KO and of cells deleted for the *pts* operon (PTS-KO) after 3 days incubation. (E) Pictures of wild-type colonies overexpressing EI-mCherry (EI-OE) or just mCherry (CTRL) grown in acidic M9 (pH 5.5) or in neutral M9 (pH 7) supplemented with 0.4% glucose with or without 0.1% of arabinose inducer for 72 hours and spotted after serial dilutions. (F) Pictures of wild-type (WT) and TmaR-KO (KO) cells grown in acidic M9 medium (pH 5.5) supplemented with the indicated carbon sources for 72 hours and spotted after serial dilutions as in (A). (G) Pictures of wild-type (WT) and TmaR-KO (KO) cells grown in acidic M9 medium (pH 5.5) supplemented with the indicated carbon sources for one week and spotted after serial dilutions.

Because fast transport of glucose is expected to lead to an increase in overflow metabolism to acetate, which acidifies the medium, we wondered if this is the cause for the decreased survival of the TmaR-KO cells. We therefore measured the pH of several wild-type and TmaR-KO cultures after growth for 3 days at pH 5.5 and observed a small, but highly consistent and reproducible difference: while the pH in the WT cultures was pH 4.1, that of the TmaR-KO was 4. Hence, the improved glucose uptake by TmaR-KO acidifies the medium faster, compared to WT, and might partly explain the difference in survival.

Monitoring cell survival in pH 5.5 in the presence of 0.4% glucose at specific time points showed that there was no significant difference in cell survival between the wild-type and TmaR-KO in the first 48 hours of growth (see SI Appendix, **Fig. S22**). Of note, reduction in cell survival in this medium was also observed for wild-type cells, but only after a week, suggesting that a decrease in cell survival at acidic pH occurs in wild-type, but the absence of TmaR dramatically increases its rate. The results thus far indicate that TmaR-mediated control of glucose consumption is important for long-term survival of *E. coli* in mild acidic conditions.

To learn more about the survivals phenotype, we viewed the wild-type and TmaR-KO cells that were grown for 3 days in minimal medium, pH 5.5, supplemented with 0.4% glucose under the microscope. One might have expected the non-growing TmaR-KO cells to undergo lysis, but instead, we observed a most distinct difference between the two strains morphology (**Fig. 6C)**. While the WT cells were small and a bit rounded, a typical morphology of *E. coli* cells in deep SP, the TmaR-KO cells were elongated, resembling more cells in logarithmic than in SP, a surprising morphology for cells that cannot grow on plates, that is, cannot divide successfully. Another difference was the detection of a bright spot near one pole of the TmaR-KO cells, which might represent protein inclusion bodies, lipids inclusions or another dense aggregation. Furthermore, when grown on fructose-supplemented MacConkey plate, the wild-type colonies turned from red to white already after one or two days, while the TmaR-KO colonies remained red even after 3 days (**Fig. 6D**) and longer (not shown), when the medium was very acidic due to sugar fermentation. Hence, TmaR-KO do not cease consuming sugar in acidic pH, although their survival in these conditions is greatly reduced compared to wild-type cells. Bacteria picked from this plate after 2 or 3 days showed the same difference in morphology observed between wild-type and TmaR-KO cells grown in liquid medium (see SI Appendix, **Fig. S23**). Hence, it seems that the TmaR-KO cells cannot enter SP properly in acidic conditions and accumulate defective molecules, leading to their death in mildly acidic conditions.

The above results raised the question whether the level of the RNA polymerase sigma factor RpoS, the primary regulator of SP genes, changes in TmaR-KO cells. Hence, we compared the level of RpoS in wild-type and TmaR-KO cells grown in acidic or neutral medium for 1, 2 or 3 days by Western blot analysis, using anti-RpoS antibodies. The results (see SI Appendix, **Fig. S24)** show that, whereas in neutral pH the level of RpoS increases gradually over time in both strains, in acidic pH, the level of RpoS in wild-type cells decreases after a day, while its level in TmaR-KO cells does not seem to change over time. Hence, reduction in TmaR-KO cells survival in acidic conditions correlates with a higher level of RpoS compared to wild-type cells, maybe hinting at an attempt to enter SP.

Since we observed that absence of TmaR leads to an increase in the kinetics of PTS sugars uptake, we asked whether the concentration of glucose in the acidic medium plays a role in TmaR-dependent cell survival. Our results show that when the glucose concentration is reduced from 0.4% to 0.1% or to 0.04%, the reduction in TmaR-KO cell survival was less severe. That is, in 0.1% glucose, the survival of the TmaR-KO cells was somewhat less reduced than in 0.4% glucose, and no difference was observed between the survivals of the two strains in 0.04% glucose (**Fig. 6A**). Increasing the glucose concentration from 0.4% to 1% did not make the survival defect of TmaR-KO more severe (**Fig. 6A**), suggesting that the cells reached saturation in glucose consumption and/or that sugar toxicity is not involved in this phenomenon. Indeed, we observed no elevation in the level of YigL, one of the hallmarks of sugar toxicity, whose level has previously been shown to elevate in sugar stress (*36, 37*), in TmaR-KO compared to WT cells grown at pH 5.5 in the presence of 0.4% glucose after 3 and 4 days (see SI Appendix, **Fig. S25**). Hence, the survival defect of Tma-KO cells in acidic conditions, although influenced by glucose concentration, is not due to sugar toxicity.

Next, we asked if EI hyperactivity for sugar consumption in TmaR-KO cells is the cause for the reduced survival in acidic pH. To address this question, we expressed EI-mCherry (EI-OE) or only mCherry (CTRL) from an arabinose-induced promoter in TmaR-KO cells and grew them in minimal medium at pH 5.5 with 0.4% glucose for 3 days. The TmaR-KO cells overexpressing EI-mCherry (0.1% arabinose) did not survive at all in these mildly acidic conditions as opposed to TmaR-KO cells overexpressing only the mCherry tag (**Fig. 6E**, middle panel). This inability of the TmaR-KO cells to survive was a direct consequence of EI overexpression, since when EI expression was not induced (no arabinose), no survival difference was observed between the strains (**Fig. 6E**, upper panel). Moreover, when growing these two strains at pH 7 with the arabinose inducer, survival of both strains was the same (**Fig. 6E**, lower panel). These results directly connect between EI activity and survival in acidic pH and suggest that EI hyperactivity in TmaR-KO cells is the mechanism that negatively affects their survival.

To understand whether the nature of the carbon source is important for the observed TmaR-dependent survival, we tested cell survival in the presence of PTS sugars other than glucose (fructose, trehalose or mannitol), whose uptake is directly affected by EI the same as glucose, of a non-PTS sugar (lactose), whose uptake is indirectly affected by EI (*38*), and of non-sugar carbon sources (casamino acids (CAA) or succinate) that are not known to be affected by EI and are incorporated into the bacterial carbon metabolism differently than sugars (*39*). When cells were grown in the presence of fructose and trehalose, TmaR-KO displayed a reduction in cell survival at 72 hours, similar to that observed with glucose; in the presence of mannitol, the reduction in cell survival was less severe; and in the presence lactose, CAA, and succinate, there was no change in survival compared to wild-type (**Fig. 6F**). However, after a week of growth, a severe defect in TmaR-KO survival was observed with mannitol and lactose, but not with CAA and succinate (**Fig. 6G**), suggesting that a drop in survival occurs in the presence of carbon sources that can enter glycolysis - PTS and non-PTS sugars – albeit at different rates, but not with non-sugar carbon sources. Taken together, our results suggest that TmaR-mediated regulation of sugar consumption (PTS or non PTS) is important for cell survival in acidic environments.

Finally, to figure out if the inability of TmaR to get phosphorylated on Y72 also affects survival in acidic conditions, we compared survival in pH 5.5 of cells expressing TmaR Y72F, TmaR Y79F and TmaR triple mutant to that of cells expressing wild-type TmaR and TmaR-KO cells. The results show that, unlike TmaR deletion, impairment of TmaR phosphorylation had no effect on survival in acidic conditions(see SI Appendix, **Fig. S26)**, in line with our finding that TmaR deletion and TmaR inability to get phosphorylated had opposite effects on sugar consumption.

## DISCUSSION

How biochemical reactions are compartmentalized within a bacterial cell is largely an open question. Unlike eukaryotes, which have organelles that enable them to control their enzymatic functions within a restricted environment, most prokaryotes do not have membrane-bounded organelles. Obviously, bacteria have developed strategies to efficiently process biochemical reactions in their crowded cytoplasm, but their nature is mostly unknown. Here, we describe a hitherto unrecognized layer of regulation of sugar metabolism in bacteria, which is based on the spatiotemporal organization of the genera PTS protein EI. We discovered TmaR, a novel protein, which localizes to the pole depending on its phosphorylation on a tyrosine residue, and show that it regulates sugar metabolism by controlling the activity of EI via polar sequestration and release, thus enabling survival in mild acidic conditions (schematically illustrated in **Fig. 7**).

**Figure 7.**
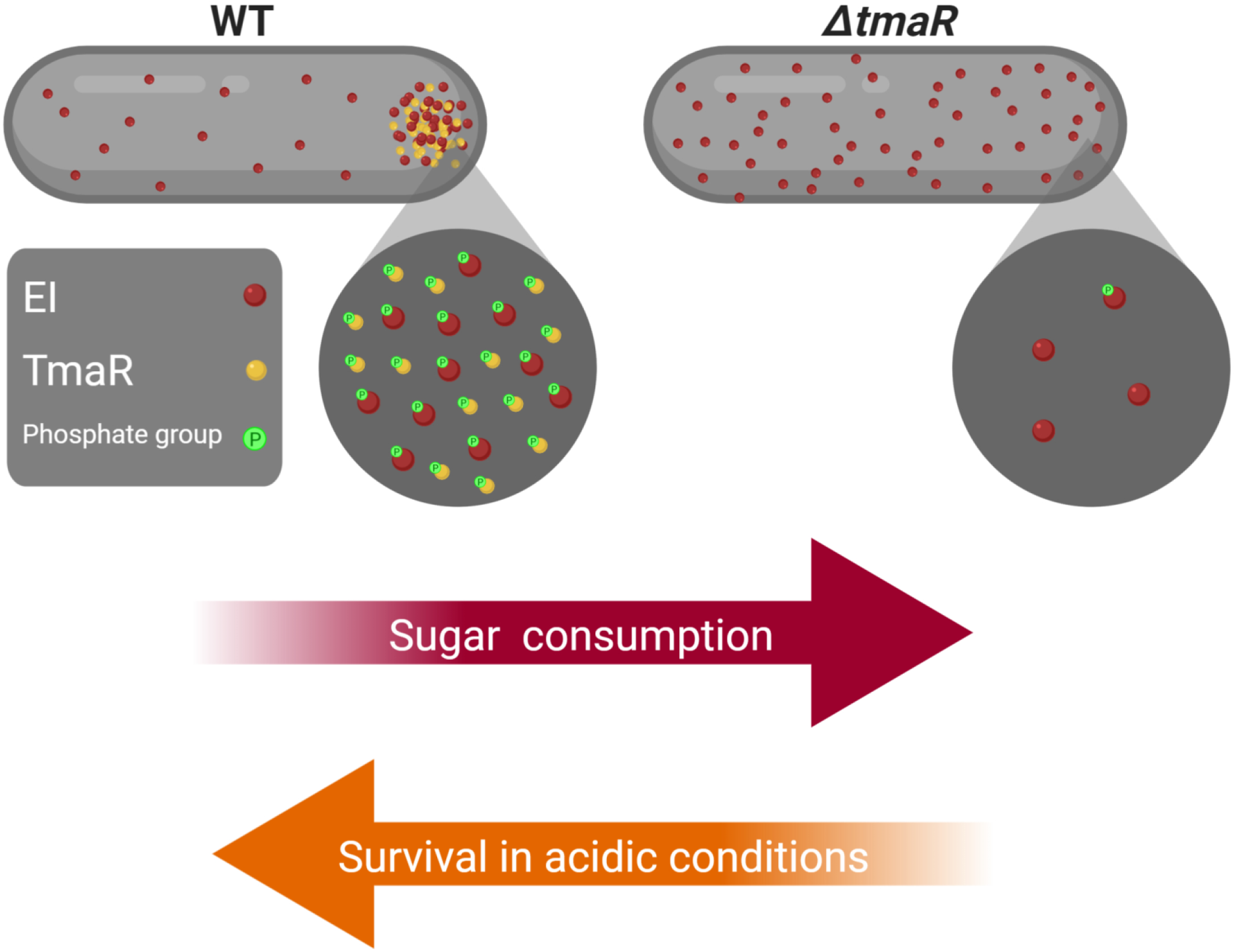
A model for TmaR-mediated control of EI localization and its implications. In wild-type cells (WT), tyrosine-phosphorylated TmaR sequesters a fraction of EI protein, which is tyrosine-phosphorylated, in polar clusters, thus restricting EI activity and enabling survival in mild acidic conditions. In Δ*tmaR* cells, EI is completely diffused, enabling hyper-consumption of sugar that is detrimental to bacterial cell survival in acidic conditions.

TmaR is the first *E. coli* protein reported to localize to the poles depending on its phosphorylation on a tyrosine residue. A SILAC-based analysis of the *E. coli* phosphoproteome predicted that TmaR is phosphorylated on TyR 79 (*21*), rather than on Tyr 72, as indicated by our results. A possible explanation for this discrepancy is that the proximity of the two residues enables phosphorylation of both, which depends on Tyr 72 being phosphorylation first. Indeed, a careful look at TmaR Y79F probed with anti-pTyr (**Fig. 1G)** reveals a slight reduction in the intensity of the band compared to WT. Phosphorylation on multiple residues, depending on the conditions has often been observed in eukaryotes (e.g., (*40*) (*41*) (*42*)). An alternative explanation is that TmaR is phosphorylated on different tyrosine residues depending on the growth phase and the environmental conditions. Indeed, phosphorylation on Tyr79 was detected during SP (*21*), whereas our results show that less TmaR molecules localize to the poles in this phase. Remarkably, all three tyrosines residues in TmaR are preceded by an aspartic acid, culminating in a DY motif, which is one of 13 statistically significant sequence motifs predicted for tyrosine phosphorylation sites in human proteins (*27*). Analysis of 512 tyrosine phosphorylation sites in *E. coli* showed underrepresentation of lysine residues in position +2 (*27*), which is the case for TyR 55, making Tyr72 and Tyr79 more favorable sites for phosphorylation. Intriguingly, the sequence around Tyr 79 matches one of the five motifs, YXXK, identified for bacterial proteins phosphorylated on tyrosine (*27*). Future research will hopefully elucidate the role of Tyr79 phosphorylation.

The kinase that phosphorylates TmaR is not known. Despite of the numerous proteins predicted to be phosphorylated on tyrosines in *E. coli*, only two BY kinases, Wzc and Etk, are known in this organism, and they both contain two transmembrane domains and are involved in cell envelope maintenance (*25, 26*). The observation that deletion of both *wzc* and *etk* genes did not diminish the extent of tyrosine phosphorylation in *E. coli* cells (*27*) indicates that additional tyrosine kinases await to be discovered in this organism.

We have previously suggested that the poles serve as hubs for sensory systems, thus enabling their communication and the generation of optimal responses (*16*). We have recently shown that RNAs that are involved in sensing and responding also accumulate at the poles (*5*). TmaR-mediated sequestration of EI molecules at the pole, highlight the importance of poles as depots that enable the production of key factors before they are needed and their release upon need, providing an additional level of regulation of sugar consumption. Such storage mechanisms have been previously described in *E. coli* for MurG and FtsZ (*8, 9*), but the proteins contributing to their polar deposition are not known. Regulation of the sugar regulator Mlc by the glucose permease PtsG has been shown to occur via membrane sequestration (*43*). Of note, tyosine phosphorylation-dependent recruitment of pole localizers in other bacteria has not been reported, although phosphorylation of DivIVA on a serine in *B. subtilis, (44*) and on a threonine in *Streptococcus suis* (*45*) were detected, but direct evidence for their occurrence *in vivo* has not been obtained yet, let alone for their involvement in polar localization.

TmaR is usually observed in one cell pole, unless overexpressed. Of note, the two poles in organisms that divide symmetrically, such as *E. coli* and *B. subtilis*, are different, one old and the other new. Various events which may potentially occur in both poles, end up taking place only in one, such as cell division during sporulation of *B. subtilis* (*46*) or plasmolysis in *E. coli* (*47*). Polar proteins often localize to one pole in these organisms, when expressed from the chromosome, although their overexpression reveals that the two poles have capacity to retain them. Other proteins were shown to oscillate from pole to pole or to relocate to midcell, which is the pole to be (*15, 48*). Future experiments with the microfluidic mother machine (*49*) are required to determine which proteins have a preference for a certain pole, old or new.

Our data showing that TmaR and EI interact *in vivo* and *in vitro* provides a mechanism for TmaR-mediated polar clustering of EI. This concept is indirectly supported by our findings that when TmaR is overexpressed, more EI molecules accumulate at the pole, whereas when EI is overexpressed, less TmaR molecules localize at the pole. Thus, assembly of the two proteins at the pole is controlled by a stoichiometric balance between them. This stoichiometry does not reflect the ratio between the total number of molecules of each of these proteins in the cell and can, therefore, be affected by changing the level of one of them. Changing the balance affects sugar consumption, as evidenced by the increased sugar consumption when TmaR is absent and EI is, therefore, non-clustered. This stoichiometric balance suggests that rather than functioning as an ON-OFF switch, TmaR functions as a fine-tuner that regulates EI activity by controlling the dynamics of EI clustering and release. For such a mechanism to be efficient, the interactions between the two proteins need to be weak and easily reversible (typical of phase separation, see below). Our results suggest that this type of interaction depends on TmaR phosphorylation on Tyr 72, since substitution of this residue to a non-phosphorylatable residue renders the interaction quite irreversible and prevents EI release from the pole. To the best of our knowledge, this is the first report of a protein, which is not a part of the PTS family that has a global effect on the PTS pathway by spatiotemporally controlling its major regulator.

We previously suggested that factors mediating EI polar clustering localize to membrane regions with strong negative curvature (*14*). However, sequence analysis does not predict any membrane-traversing or -binding motifs in TmaR. Additionally, our previous (*15*) and present results show that both EI and TmaR are dynamic, mainly within the pole region and do not appear to be membrane-anchored, suggesting that their localization involves additional mechanisms, such as the formation of condensates via liquid-liquid phase separation (LLPS) (*50*). Evidence supporting this notion is the FRAP analysis of EI that shows the recruitment of EI molecules into the cluster in few minutes (*15*), which might indicate high fluidity of the cluster, characteristic of LLPS (*51*). Moreover, both TmaR and EI are predicted to have unstructured domains (*52*), a common feature of proteins capable of LLPS (*51*). Also tyrosine phosphorylation, shown here to be important for TmaR and EI localization, has been implicated in LLPS in eukaryotes, as it enables electrostatic interactions that mediate LLPS (*53*). Because TmaR controls EI localization, but not vice versa, TmaR is expected to be the one to induce the putative LLPS. Our finding that TmaR limits the degree of heterogeneity in EI activity is in line with this hypothesis, since compartmentalization of proteins by LLPS has been suggested as a mechanism to reduce noise (*54*). Further research to examine this possibility is required.

The phosphorylated tyrosine residue in EI, Tyr122, is a conserved residue between EI^sugar^ and EI^Ntr^ (*33*), suggestive of its importance. Notably, extensive tyrosine phosphorylation occurs in proteins linked to sugar metabolism and the TCA cycle in *E. coli* and in the archaeon *Sulfolobus solfataricus* (*55*), suggesting that the impact of tyrosine phosphorylation on proteins that function in carbohydrate metabolism is conserved through evolution. The finding that tyrosine phosphorylation regulates the activity of the catabolite regulator Cra (*22*), together with our findings that it regulates localization of EI and of the factor that controls EI activity, TmaR, supports this hypothesis. The finding that among the proteins that co-purified with TmaR, the biggest group was of proteins that are involved in metabolic processes strengthen this assumption.

The multifaceted effect of TmaR on regulating sugar metabolism and cell survival at low pH explains why TmaR is a conserved protein. Although TmaR is not essential in laboratory growth conditions, it is apparently essential for bacterial survival in their natural habitats, e.g., the human colon (pH 6-7.5), the caecum, which is the beginning of the large intestine (pH 5.7) and the small intestine (pH 4-6) (*56*), as well as macrophages (pH ~5.0) (*34*). As for sugars, glucose concentration in the human colon is estimated to reach up to 40 mM (*57*), but food intake and digestion lead to rapid changes in gut sugar concentrations. Hence, counteracting the stress associated with high sugar concentration and low pH is particularly essential in the gut. We suggest that TmaR makes an important regulatory connection between sugar uptake and cell survival in mild acidic conditions, since survival of TmaR-KO cells under these conditions is observed only when the cells are grown with sugars and not with other carbon sources. This might be explained by the production of lactate or pyruvate and/or due to EI phosphorylation by PEP that produces pyruvate, both due to the sugars that enter glycolysis (*59*). Overproduction of pyruvate or its acidic derivatives can lower the pH and create harmful conditions that eventually might kill the cells and, therefore, require acid resistance (*57*). Importantly, overexpression of EI reduces survival in mildly acidic conditions just like deletion of TmaR, strengthening our assumption that this effect is mediated through EI hyperactivity. Because EI and the PTS in general play roles in many regulatory pathways, such as carbon catabolite repression, nitrogen and phosphate metabolism, potassium transport and chemotaxis (*58, 59*), the effect of TmaR on EI activity is expected to have a far bigger effect on cell physiology than just the control of sugar flux.

Why do cells lacking TmaR die in mild acidic pH? Our results argue against the possibility that they are more susceptible to sugar toxicity. On the other hand, the morphology of the TmaR-KO cells after growing them for 3 days at pH 5.5 suggest that they cannot properly enter SP, a phase that enables cells to adapt to and survive in suboptimal environments that involve nutrient deprivation. Entry into SP requires complex and coordinated changes in gene expression, which enable reduction in cell metabolism and energy conservation. Because bacteria spend most of their time in nature in stressful conditions and intense competition for resources, difficulties in entering SP is a severe defect that makes cells waste their limited resources, much like “let us eat and drink, for tomorrow we shall die” (Isaiah 22). This is exemplified by the red TmaR-KO colonies after 3 days on MacConkey plates, as opposed to wild-type colonies that turn completely white.

The alternative sigma factor RpoS controls the SP-related changes in gene expression, allowing cells to become more resistant not only to the stress that they first encounter but also to other stressful treatments (*60, 61*). The report on the increase in RpoS levels in various stresses (*62*), which reflects the need to combat with stressful conditions might explain the elevation in RpoS level in TmaR-KO cells compared to wild-type in acidic conditions. The linkage between the control of *rpoS* expression and carbon metabolism suggests that the uncontrolled activity of EI in TmaR-KO cells affects proper entry into SP. Of note, expression of *tmaR* is dramatically increased in Δ*rpoS* cells (*63*), suggesting that control of TmaR level is important for survival in SP.

## MATERIALS AND METHODS

All methods and materials used for this study are described in the Supporting Information. This includes constructions of all strains, fluorescence microscopy and image analysis, biochemical methodologies used for protein analysis and protein-protein interaction and microbial practices.

## Supporting information

see SI Appendix

## ACKNOWLEDGMENTS

We thank Avital Cher for screening the YFP-tagged ORFs for their localization, Omer Goldberger for help with the illustration and Mikel Irastorza for critical reading of the manuscript. We acknowledge Joachim O. Radler and Abram Aertsen for the gift of T1683 and IbpA-msfGFP strains, respectively. We are grateful to Susan Gottesman, Teppei Morita and Hiroji Aiba for the generous gift of α–RpoS and α IIEB^Glc^ antibodies, as well as to Ophry Pines and Sigal Ben-Yehuda for sharing antibodies against His-tag and GFP-tag, respectively. We thank Reuven Wiener for help with the pET plasmids, Yair Katz for help with Matlab and Excel, Meshi Barsheshet for help with analysis with R, and Yair E. Gatt for generating TmaR phylogenetic tree. We thank members of Orna Amster-Choder and Hanah Margalit labs for fruitful discussions and Regine Hengge and Roberto Kolter for consultation about RpoS. This research was supported by the Israel Science Foundation (ISF) founded by the Israel Academy of Sciences and Humanities (1274/19) and the Deutsch-Israeli Project Cooperation (DIP) (AM 441/1-1 SO 568/1-1).

## Notes

### Competing Interest Statement

The authors have declared no competing interest.

