## Supplementary material for "Tyrosine phosphorylation-dependent localization of TmaR, a novel *E. coli* polar protein that controls activity of the major sugar regulator by polar sequestration": see SI Appendix

**This PDF file includes:**

Materials and Methods

Table S1 to S3

References

Figures S1-S26

### Construction of the strains and plasmids

TS-TmaR-YFP, which expresses YFP-tagged TmaR from the chromosome of BW25113, was constructed by P1 transduction (*1*) of *yeeX-YFP::cat* from SX1989 (*2*) to BW25113. TS-mYFP-TmaR, which expresses monomeric YFP fused to the N terminus of TmaR from BW25113, was constructed by three overlapping PCR amplification of: i) the sequence preceding *tmaR*, ii) *mVenus* with a short peptide linker: GGGGSGGGGSSG and iii) *tmaR* plus the sequence following it. The three amplicons were ligated using Gibson assembly (*3*). The assembled sequence was than amplified and introduced into BW25113, using positive and negative selection (*4*). TS-EI-mCherry, which expresses EI-mCherry from BW25113, was constructed by P1 transduction of *ptsI-mCherry::kanR* from MG1655ϕEImCherry (*5*) to BW25113. TS-TmaR-YFP+EI-mCherry, which expresses both TmaR-YFP and EI-mCherry from BW25113, was constructed by P1 transduction of *yeeX-YFP::cat* from SX1989 to TS-EI-mCherry. TS-TmaR-KO, which has a deletion of the *tmaR* gene in BW25113 was constructed by P1 transduction of *ΔyeeX::kan* from JW1989-1 (*6*) to BW25113. TS-TmaR-KO+EI-mCherry was constructed by first removing the $Kan^R$ cassette from TmaR-KO, using pCP20 (*7*), and then by P1 transduction of *ptsI-mCherry::kanR* from EI-mCherry to TmaR-KO with no $Kan^R$. TS-TmaR-YFP+PTS-KO (MG) was constructed by P1 transduction of *yeeX-YFP::cat* from SX1989 to Δ*pts* (*8*). TS-TmaR-YFP (MG) was constructed by P1 transduction of *yeeX-YFP::cat* from SX1989 to MG1655. NTS102-TmaR-KO was constructed by first removing the $Kan^R$ cassette from NTS102, using pCP20, and then by P1 transduction of *ΔyeeX::kan* from JW1989-1 to NTS102 with no $Kan^R$ (*9*). TS-YigL-FT was constructed by three overlapping PCR amplification of: i) the *yigL* gene with a primer encoding for 3XFlag tag, ii) $Kan^R$ with FRT sequence on both sides, and iii) the sequence following *yigL*. The three amplicons were ligated using Gibson assembly. The assembled sequence was than amplify and introduced into BW25113 by lambda red insertion with selection for kan resistance (*10*). TS-TmaR-KO+YigL-FT was constructed by first removing the $Kan^R$ using pCP20, and then by P1 transduction of *ΔyeeX::kan* from JW1989-1 to YigL-FT. TS-TmaRY79F, TS-TmaRY72F and TS-TmaRY51F were constructed by two overlapping PCR amplifications of the *tmaR* gene with the sequences preceding and following it and with primers containing the respective mutations, which were ligated using Gibson assembly. The assembled sequence was than amplified and introduced into BW25113 using positive and negative selection. The double and triple mutations TS-TmaRY79F,Y72F and TS-TmaRY79F,Y72F,Y51F were constructed in a similar way using TS-TmaRY79F and TS-TmaRY79F,Y72F respectively as templates for the PCR. TS-TmaRY79F-YFP, TS-TmaRY72F-YFP, TS-TmaRY51F-YFP, TS-TmaRY79F,Y72F-YFP and TS-TmaRY79F,Y72F,Y51F-YFP were constructed by two overlapping PCR amplifications of: i) TS-TmaRY79F, TS-TmaRY72F, TS-TmaRY51F, TS-TmaRY79F,Y72F or TS-TmaRY79F,Y72F,Y51F, respectively, and ii) TS-TmaR-YFP. The two fragments were combined by Gibson assembly. The assembled sequence was than amplifid and introduced into BW25113 using positive and negative selection. TS-mYFP-TmaRY79F, TS-mYFP-TmaRY72F,

TS-mYFP-TmaRY51F, TS-mYFP-TmaRY79F,Y72F and TS-mYFP-TmaRY79F,Y72F,Y51F were constructed by two overlapping PCR amplifications of: i) TS-mYFP-TmaR and ii) TS-TmaRY79F, TS-TmaRY72F, TS-TmaRY51F, TS-TmaRY79F,Y72F or TS-TmaRY79F,Y72F,Y51F, respectively. The two fragments were combined by Gibson assembly. The assembled sequence was than amplified and introduced into BW25113 using positive and negative selection. TS-TmaRY72D and TS-TmaRY72E were constructed by two overlapping PCR amplifications of the *mVenus-tmaR tmaR* gene with the sequences preceding and following it and with primers containing the respective mutations, which were ligated using Gibson assembly. The assembled sequence was than amplified and introduced into BW25113 using positive and negative selection. TS-TmaR-YFP+Wzc-KO was constructed by P1 transduction of *ΔwzC::kan* from JW2045 (*6*) to TS-TmaR-YFP. TS-TmaR-YFP+Etk-KO was constructed by P1 transduction of *ΔyccC::kan* from JW0964 (*6*) to TS-TmaR-YFP. TS-BL21+EI-mCherry(no-kan$^R$) was constructed by transducing *ptsI-mCherry::kanR* from MG1655ϕEImCherry (*5*) to BL21(DE3) via P1 and then removing the Kan$^R$ cassette using pCP20. TS-BL21(DE3)+EI-mCherry+IbpA-msfGFP was constructed by transducing *IbpA-msfGFP::kan* from IbpA-msfGFP strain (*11*) to TS-BL21+EI-mCherry(no-kan$^R$) via P1. TS-EIY122F-mCherry was constructed by two overlapping PCR amplifications of *ptsI-mCherry::kanR* with the sequences preceding and following it and with primers containing the respective mutations, which were ligated using Gibson assembly. The assembled sequence was than amplified and introduced, using lambda red insertion, into *ΔptsI::kan* from JW2409 (*6*), from which the Kan$^R$ cassette has been removed by pCP20. After the insertion, we selected for Kan resistance. TS-TmaR-YFP+EI Y122F-mCherry was constructed by P1 transduction of *ptsI Y122F-mCherry::kanR* from TS-EIY122F-mCherry to TS-TmaR-YFP. TS-EI-mCherry+Wzc-KO was constructed by first removing the Kan$^R$ cassette from TS-EI-mCherry, using pCP20, then using P1 to transduce *ΔwzC::kan* from JW2045 (*6*) to TS-EI-mCherry without Kan$^R$. Similarly, TS-EI-mCherry+Ezc-KO was constructed by first removing the Kan$^R$ cassette from TS-EI-mCherry, using pCP20 and then transducing *ΔyccC::kan* from JW0964 (*6*) to TS-EI-mCherry without Kan$^R$ by P1.

We constructed pET15b-TmaR using two overlapping PCR products that were combined by Gibson assembly. The first amplicon was of the pET15b plasmid (Addgene Plasmid 29653) using the following primers: F-CTCGAGGATCCGGCTGCTAAC and R-CCCCTGAAAGTAAAGATTCTCC. The second amplicon was of the *tmaR* gene amplified from BW25113 using the following primers: F-TATGGAGAATCTTTACTTTCAGGGGatgGAAACTACCAAGCCTTC and R-GGCTTTGTTAGCAGCCGGATCCTCGAGttaCTTCGCTTCGCCG. For pET15b-TmaRY79F, pET15b-TmaRY72F, pET15b-TmaRY51F, pET15b-TmaRY79F,Y72F, pET15b-TmaRY79F,Y72F,Y51F we used two overlapping PCR products that were later attached by Gibson assembly. The first PCR, which was similar to all five plasmids, was of pET15b with the N and the C termini of the *tmaR* gene amplified from pET15b-TmaR with the following primers: F-

CCGCGATATCTCCAAAAAGCTG and R-CAGTTTGTTCTTACGACGGAAC. For the second PCR, *tmaR* Y79F, *tmaR* Y72F, *tmaR* Y51F, *tmaR* Y79F,Y72F or *tmaR* Y79F,Y72F,Y51F was amlified from the chromosome of cells carrying the respective mutation, using the following primers: F-GTTCCGTCGTAAGAACAAACTG and R-CAGCTTTTTGGAGATATCGCGG. pBAD18-mCherry was constructed by amplifying mCherry from pBADLLEI-mCherry using the following primers: F-GCGGCTAGCCAATTCCCCTCTAGAAATAATTTTGTTTAACTTTAAGAAGGAGATATACCATGGTGAGCAAGGG and R- ATGGTCGACTTACTTGTACAGCTCG. The forward primer contained a ribosomal binding site from pET-15b (CTAGCCAATTCCCCTCTAGAAATAATTTTGTTTAACTTTAAGAAGGAGATATACC) to have the same control elements as pBADLLEI-mCherry. The PCR product was cleaved with NheI and SalI (added to the primers) and then ligated to pBADLLEI-mCherry, which was also digested with NheI and SalI. To construct pCA24N-TmaRY79F, pCA24N-TmaRY51F, pCA24N-TmaRY79F,Y72F and pCA24N-TmaR Y79F,Y72F,Y51F, we used two overlapping PCR products that were later attached by Gibson assembly in the same way and same primers as described above for pET15b plasmids with one modification: the template for the first PCR was JW1989-*yeeX* (pCA24N). pBADLLEI Y122F-mCherry was constructed by amplifying *ptsI* from TS-EIY122F-mCherry cells, using F-GCATCCCCGGGTATCGCTTTCG and R- CTACCGGTACCCACGATAGC primers. The PCR product and pBADLLEI-mCherry were then cleaved with the restriction enzymes XmaI and KpnI and ligated. We constructed pET-GFP-TmaR using two overlapping PCR products that were combined by Gibson assembly. The first amplicon was of the pET-GFP plasmid (Addgene Plasmid 29663) using the following primers: F- TAACGGATCCGAATTCGAGC and R-TCCACTTCCAATATTGGATTGG. The second amplicon was of the *tmaR* gene amplified from BW25113 using the following primers: F-CTTCCAATCCAATATTGGAAGTGGAatgGAAACTACCAAGCCTTC and R-ACGGCGCTCGAATTCGGATCCGttaCTTCGCTTCGCCG. To construct pET-GFP-TmaRY72F, we used two overlapping PCR products that were later attached by Gibson assembly in the same way and same primers as described above for pET15b-TmaR Y72F plasmid, with one modification: the template for the first PCR was pET-GFP-TmaR.

**Growth conditions**

Unless otherwise indicated, overnight cultures were grown at 37 $^{o}$C in LB and diluted 1:100 into LB supplemented with the appropriate antibiotics at the following concentrations: kanamycin (30 µg/ml), chloramphenicol (25 µg/ml) and ampicillin (200 µg/ml). If not indicated that the protein was overexpressed, it was expressed from its gene's native promoter and locus in the chromosome. Unless otherwise indicated, expression from plasmids was induced by 0.1 mM IPTG or 0.1% arabinose, depending on the promoter, from the start of growth. Unless otherwise indicated, growth was continued until mid-logarithmic phase.

**Fluorescence microscopy**

Fluorescence microscopy was carried out as described previously (*9*). For snap-shot imaging, 0.7 ml of cells were centrifuged, re-suspended in 10–300 µl (according to the cell density) of fresh LB and spotted on 1% agarose LB pads on a slide. Time-lapse imaging was performed, as previously described (*9*). Unless otherwise indicated, the scale bar size was 5 µm. Cells are shown in phase (grey), in the appropriate florescent channel (colored) or as merged.

**Image analysis**

Image analysis was performed with NIS-Elements Advanced Research (AR) version 4.4 software (Nikon). The detection of clusters was calculated manually using NIS Elements AR. Unless otherwise indicated, for graphs we used GraphPad Prism version 6.00 or Windows, GraphPad Software, San Diego, California USA, https://www.graphpad.com/. In Fig. 2B, the fluorescent signal along the cell axis was calculated manually using NIS Elements AR, and a heat map was then created using Microsoft Excel. In Fig. 2D, after manual cluster detection using NIS Elements AR, the colors in every row were generated using Microsoft Excel. The Nag promotor activity was analyzed using Microsoft Excel. The fluorescence intensity profile in Fig. 1B and the box plot in Fig. S15 were created using MATLAB 2017b, The MathWorks, Inc., Natick, Massachusetts, United States.

**Western Blot Analysis**

Equal amount of cells grown to the indicated $OD_{600}$ were collected from each strain, washed with PBS and laemmli buffer was added to them. Each sample was then heated to 95 $^{o}$C for 10 minutes and the lysates were separated on 12% SDS–polyacrylamide gels (unless otherwise indicated). Gels were subjected to Western blot analysis as described previously (*12*). When indicated, the nitrocellulose membrane was stained with Ponceau S (Sigma-Aldrich) and then probed with the indicated antibodies. For quantification of the Western blot bands intensity in figures 5Ac, S3 and S10 we used image J (*13*). When indicated, gels that were run in parallel were stained with Coomassie Brilliant Blue (Expedeon InstantBlue® Protein Stain).

**Far-western analysis**

Far Western analyses were carried out essentially as described previously (*14*). For Figures 2E and S12, overnight cultures of Δ*tmaR* cells overexpressing TmaR or only a His tag from pCA24N-TmaR or pCA24N, respectively, were diluted 1:100 in 40 ml of fresh LB supplemented with 0.1 mM IPTG and grown until mid-logarithmic phase. Subsequently, cells were pelleted, washed in 1X PBS and lysed by Mixer Mill MM400 using glass beads. The lysates were then mixed with laemmli buffer, boiled at 95 $^{o}$C for 10 minutes and fractionated on a 12% SDS polyacrylamide gel. The proteins were blotted onto a nitrocellulose two membranes. The first membrane (for Western analysis) was incubated overnight in 4 $^{o}$C with anti-His antibodies and then anti-mouse antibodies The second membrane (for Far-Western analysis) was incubated with purified EI-mCherry or mCherry in 7% Difco skim milk (BD biosciences) diluted in 10 ml PBST (Phosphate buffered saline with Tween-20) for overnight in 4 $^{o}$C, washed, incubated with anti-mCherry antibody for overnight in 4 $^{o}$C, washed trice and incubated with anti-rabbit antibody for one hour. For purification of EI-mCherry or mCherry, overnight cultures of BW25113 cells with pBADLLEI-mCherry or pBAD18-mCherry were diluted 1:100 in 40 ml of fresh LB supplemented with 0.1% arabinose, grown until mid-log, lysed by Mixer Mill MM400 using glass beads, purified using RFP-Trap®_A (Chromotek), as suggested by the manufacturer, and eluted by adding 50 µl 0.1 M glycine at pH 2.5 (incubation time: 30 sec – 2 min) followed by neutralization with 5 µl of 1 M Tris-base, as suggested by the manufacturer, to preserve the native structure of EI-mCherry or mCherry.

For Fig. 5A and 5F, overnight cultures of Δ*tmaR* cells overexpressing TmaR or its variants (TmaR Y79F, TmaR Y51F, TmaR Y79F+Y72F or TmaR Y79F+Y72F+Y51F) from the respective PCA24N plasmid derivatives (PCA24N-TmaRY79F, etc., see Table S1), as well as only a His tag from PCA24N were diluted 1:100 in 5 ml of fresh LB supplemented with 0.1 mM IPTG and grown until mid-logarithmic phase. Subsequently, 0.5 ml cells were pelleted, washed with 1X PBS, re-suspended in laemmli buffer, heated to 95 $^{o}$C for 10 minutes and fractionated on 4-20% gradient SDS polyacrylamide gel. The proteins were blotted onto a nitrocellulose membrane that was probed with EI-mCherry (Fig. 5A) or EI Y122F-mCherry (Fig. 5F) followed by anti-mCherry antibodies, for two hours washed trice and subjected to an anti-rabbit antibody for one hour. After exposing the membrane using ECL, the membrane was stripped and incubated with anti-His tag2 antibodies overnight in 4 $^{o}$C followed by anti-mouse mouse antibodies for 1 hour.

For the purification of EI-mCherry and EI Y122F-mCherry, overnight cultures of MG1655 cells carrying pBADLLEI-mCherry or pBADLLEIY122F-mCherry were diluted 1:100 in 50 ml of fresh LB supplemented with 0.1% arabinose and grown until mid-logarithmic phase. Subsequently, cells were pelleted, washed in 1X PBS with 10 mM imidazole, and 1 mM AEBSF and lysed by Mixer Mill MM400 using glass beads. Purification was done as recommended by the manufacture (Thermo Fisher Scientific HisPur™ Ni-NTA Resin) with minor changes: the mixing of the resin and the lysate were done for 2h, the incubation of the elution buffer and the resin was for 15 minutes. The total cell lysate, the flow-through and the elations were later fractionated by SDS-PAGE and stained with Coomassie Brilliant Blue.

**MacConkey plates**

MacConkey plates were made by mixing 40 mg/ml of Difco MacConkey Agar base (Becton Dickinson and company) with 0.4% Fructose. Bacteria were streaked from fresh single colonies, and the plates were incubated overnight at room temperature, unless otherwise indicated. Cells deleted for the *pts* operon (PTS-KO) served as a negative control (growth of white colonies) for consumption of PTS sugars on the MacConkey plates. For Fig. S23, we picked the cells from the MacConkey plate suspended them in 30 µl PBS and observed them under the microscope**.**

**Glucose consumption assay**

This protocol was performed as previously described (*15*) with minor changes. Briefly, overnight cultures of BW25113 (WT), TmaR-KO, TmaR-OE, and Δ*pts* cells were diluted 1:100 in 50 ml LB supplemented with 25 mM glucose and 0.1 mM IPTG and grown until $OD_{600}$=0.6-0.8. Cells were collected and washed thrice with 10 ml of phosphate-buffered saline (PBS). Subsequently, pellets are re-suspended in PBS to an $OD_{600}$ of 1.5. Four ml of the suspension were taken for pre-incubation at 37°C for 5-10 minutes. Glucose was added to a final concentration of 1 mM and 300 µl of the sample were immediately withdrawn (Time 0). Samples of 300 µl were withdrawn every 5 minutes for 45 minutes. Each sample was boiled for 10 minutes at 100°C and then centrifuged at 10,000 RPM for 3 minutes. Two hundred and fifty microliters of the supernatant were mixed with 250µl of PBS and 500µl of the glucose assay reagent (Sigma G3293-50ML). After incubating for 15 minutes at room temperature, the absorbance of the reaction mixture at 340 nm was recorded. The glucose concentration was then measured by comparison to a calibration curve of different glucose concentrations. The glucose consumption rate was calculated from the slope of the calibration curve.

**NAG (N-acetyl glucosamine) promotor activity**

These measurements were performed as previously described (*9*). To measure the correlation between *pNAG-mCerulean* expressions to EI-mCherry distribution, the reporter strains were grown overnight in M63 minimal media (*16*) supplemented with 0.2% Casamino acids, 1 mM $MgSO_4$, 2 µg/ml vitamin B1 and 0.4% glycerol (M63-glycerol). Cultures were then diluted 1:100 in fresh M63-glycerol medium, grown until $OD_{600=}$0.3-0.35, washed and re-suspended in M63 supplemented with vitamin B1, $MgSO_4$ and 0.4% N-acetyl glucosamine (NAG). Snapshots of phase contrast and fluorescence signal were taken after 1 hour of incubation at 37 °C.

**Survival experiment**

Unless otherwise indicated, when cells were grown in neutral M9 medium (pH 7), commercial M9 (Sigma Aldrich) was supplemented with 2 µg/ml vitamin B1 and 1 mM $MgSO_4$. When cells were grown in acidic M9 medium (pH 5.5), a modified M9 (M9*) was prepared by mixing 42 mM $NaH_2PO_4$, 24 mM $KH_2PO_4$, 4.7 mM $Na_2HPO_4$, 1.8 mM $NH_4Cl$, 0.8 mM NaCl, 2 µg/ml vitamin B1 and 1 mM $MgSO_4$. Both media were supplemented with the indicated carbon source. For the results presented in Fig 6B, the pH of M9* was increased from 5.5 to 7 using NaOH; to lower the pH to 5 or 4, M9 was titrated with HCl. For the results presented in Fig. 6F and 6G, the carbon source was supplemented at a concentration of 0.05 M for all sugars and 0.4% for succinate and CAA. For all survival experiment, overnight cells were diluted 1:100 (time 0) in the indicated medium and grown at 37 °C. When indicated, expression of was induced with arabinose (at the indicated concentration. For plating, cells were serially diluted in PBS then plated on LB plates at the indicated dilutions (each drop contained 5 µl). After two days in RT, pictures of the plates were taken using Biorad gel doc.

**Protein analysis by mass spectrometry and data processing**

BW25113 (WT) and TmaR-YFP cells were lysed using microfluidizer and proteins were purified using GFP-Trap®_A (Chromotek), as suggested by the manufacturer. Samples were subjected to trypsin digestion and analyzed separately. Raw data was searched against a protein database of *E. coli* K12 sequences. The fold change for the comparison was calculated based on the measured protein intensity for each experiment. The volcano plot was created using R Core Team (2013) (http://www.R-project.org/). Outliers were excluded. The pie chart was created using PANTHER (*17*) in the GO annotation website (http://geneontology.org/).

**Phylogenetic tree**

We constructed a dendrogram to assess the evolutionary conservation of TmaR (YeeX). We first extracted the protein amino acid sequence from the proteome of *Escherichia coli* K12 W3110. Blastp (*18*) was then used to query the sequence against a database of 1420 complete proteomes from NCBI's RefSeq database (*19*) and compile the top hits for each genome, adhering to thresholds of *e-value* < 0.05 and identity percentage > 0.4. The hits for the different genomes were then aligned using Clustal Omega (default parameters) (*20*) to construct a multiple sequence alignment with the guide tree extracted for the dendrogram. iTOL (*21*) was used for visualization. All scripts used for this analysis are available in the following github repository: https://github.com/YairGatt/ConservationProfiler

**Schematic illustration**

The scheme in Fig. 7 was created with https://biorender.com/**.**

**Table S1. Bacterial strains used in this study and their relevant phenotype**

| **Strains** | **Relevant genotype** | **Source / Reference** |
|---|---|---|
| BW25113 (WT) | F-, *Δ(araD-araB)567*, *ΔlacZ4787*::rrnB-3, $\lambda^-$, *rph-1*, *Δ(rhaD-rhaB)568*, *hsdR514* | (*6*) |
| TS-TmaR-YFP | BW25113, *tmaR-YFP::cat* | This work |
| TS-mYFP-TmaR | BW25113, *mVenus-tmaR* | This work |
| TS-EI-mCherry | BW25113, *ptsI-mCherry::kanR* | This work |
| TS-TmaR-YFP+EI-mCherry | BW25113, *tmaR-YFP::cat*, *ptsI-mCherry::kan* | This work |
| TS-TmaR-KO | BW25113, *ΔtmaR::kan* | This work |
| TS-TmaR-KO+EI-mCherry | BW25113, *ptsI-mCherry::kanR, ΔtmaR* | This work |
| ∆pts (PTS-KO) | MG1655, *∆pts::kan* | (*8*) |
| TS-TmaR-YFP+PTS-KO(MG) | MG1655, *∆pts::kan, tmaR-YFP::cat* | This work |
| TS-TmaR-YFP(MG) | MG1655, *tmaR-YFP::cat* | This work |
| NTS102 | T1683, *ptsI-mCherry::kan, Pnag-mCerulean* | (*9*) |
| NTS102-TmaR-KO | T1683, *ptsI*-mCherry, *Pnag*-mCerulean, *ΔtmaR::kan* | This work |
| TS-YigL-FT | BW25113, *yigL-3XFT* | This work |
| TS-TmaR-KO+YigL-FT | BW25113, *yigL-FT*, *ΔtmaR::kan* | This work |
| TS-TmaRY79F | BW25113, *tmaR Y79F* | This work |
| TS-TmaRY72F | BW25113, *tmaR Y72F* | This work |
| TS-TmaRY51F | BW25113, *tmaR Y51F* | This work |
| TS-TmaRY79F,Y72F | BW25113, *tmaR Y79F, Y72F* | This work |

| | | |
|---|---|---|
| TS-TmaRY79F,Y72F,Y51F | BW25113, *tmaR Y79F, Y72F, Y51F* | This work |
| TS-TmaRY79F-YFP | BW25113, *tmaR Y79F-YFP::cat* | This work |
| TS-TmaRY72F-YFP | BW25113, *tmaR Y72F-YFP::cat* | This work |
| TS-TmaRY51F-YFP | BW25113, *tmaR Y51F-YFP::cat* | This work |
| TS-TmaRY79F,Y72F-YFP | BW25113, *tmaR Y79F, Y72F-YFP::cat* | This work |
| TS-TmaRY79F,Y72F,Y51F-YFP | BW25113, *tmaR Y79F, Y72F, Y51F-YFP::cat* | This work |
| TS-mYFP-TmaRY79F | BW25113, *mVenus-tmaR Y79F* | This work |
| TS-mYFP-TmaRY72F | BW25113, *mVenus-tmaR Y72F* | This work |
| TS-mYFP-TmaR Y51F | BW25113, *mVenus-tmaR Y51F* | This work |
| TS-mYFP-TmaRY79F,Y72F | BW25113, *mVenus-tmaR Y79F, Y72F* | This work |
| TS-mYFP-TmaRY79F,Y72F,Y51F | BW25113, *mVenus-tmaR Y79F, Y72F, Y51F* | This work |
| TS-mYFP-TmaRY72D | BW25113, *mVenus-tmaR Y72D* | This work |
| TS-mYFP-TmaRY72E | BW25113, *mVenus-tmaR Y72E* | This work |
| TS-TmaR-YFP+Wzc-KO | BW25113, *tmaR-YFP::cat*, *ΔwzC::kan* | This work |
| TS-TmaR-YFP+Etk-KO | BW25113, *tmaR-YFP::cat*, *ΔyccC::kan* | This work |
| TS-BL21(DE3)+EI-mCherry+IbpA-msfGFP | BL21(DE3), *ptsI- mCherry, ibpA-msfgfp::kan* | This work |

| | | |
|---|---|---|
| TS-BL21+EI-mCherry(no-kan$^R$) | BL21(DE3), *ptsI- mCherry* | This work |
| TS-EIY122F-mCherry | BW25113, *ptsI- Y122F mCherry::kanR* | This work |
| TS-TmaR-YFP+EIY122F-mCherry | BW25113, *tmaR-YFP::cat*, *ptsI- Y122F mCherry::kanR* | This work |
| TS-EI-mCherry+Wzc-KO | BW25113, *ptsI-mCherry::kanR*, *ΔwzC::kan* | This work |
| TS-EI-mCherry+Etk-KO | BW25113, *ptsI-mCherry::kanR*, *ΔyccC::kan* | This work |
| *ΔrpoS* | MG1655, *rpoS::Tn10* | (*22*) |

**Table S2. Plasmids used in this study and the proteins they encode**

| Plasmids | Encoded protein(s) / Inducer | Reference |
|---|---|---|
| pET15b-TmaR | TmaR/IPTG | This work |
| pET15b-TmaRY79F | TmaR Y79F/IPTG | This work |
| pET15b-TmaRY72F | TmaR Y72F/IPTG | This work |
| pET15b-TmaRY51F | TmaR Y51F/IPTG | This work |
| pET15b-TmaRY79F,Y72F | TmaR Y79F, Y72F/IPTG | This work |
| pET15b-TmaRY79F,Y72F,Y51F | TmaR Y79F, Y72F, Y51F/IPTG | This work |
| JW1989-YeeX (pCA24N-TmaR) | His -TmaR/IPTG | (*23*) |
| pCA24N | His-Tag/IPTG | (*23*) |
| pBADANSHPr-GFP | HPr-GFP/arabinose | (*5*) |
| pBADLLEI-mCherry | His-EI-mCherry/arabinose | (*5*) |
| pBAD18-mCherry | mCherry/arabinose | This work |
| pQELL-EI | His-EI/IPTG | (*24*) |
| pYT103 | PtsG-FLAG | (*25*) |

| | | |
|---|---|---|
| pCA24N-TmaRY79F | His –TmaR Y79F/IPTG | This work |
| pCA24N-TmaRY51F | His –TmaR Y751F/IPTG | This work |
| pCA24N-TmaRY79F,Y72F | His –TmaR Y79F Y72F/IPTG | This work |
| pCA24N-TmaRY79F, Y72F,Y51F | His –TmaR Y79F Y72F Y51F/IPTG | This work |
| pET-GFP-TmaR | GFP-TmaR/IPTG | This work |
| pET-GFP-TmaRY72F | GFP-TmaR Y72F/IPTG | This work |
| pBADLLEI Y122F-mCherry | His-EI Y122F-mCherry/IPTG | This work |

**Table S3. Antibodies used in this study and their source**

| antibody | source | identifier |
|---|---|---|
| α -GroEL | abcam | ab90522 |
| α -mCherry | abcam | ab167453 |
| α –His tag | Ophry Pines | N/A |
| α –His tag2 | genscript | 6G2A9 |
| α –GFP | Sigal Ben-Yehuda | N/A |
| α –GFP2 | clontech | 632592 |
| α –$IIB^{Glc}$ | Teppei Morita | N/A |
| α –FLAG | Sigma Aldrich | F1804 |
| α –phosphorylated tyrosine (4G10) | EMD MILIPORE | 05-321 |
| α – RpoS (RpoS antisera) | Susan Gottesman | N/A |

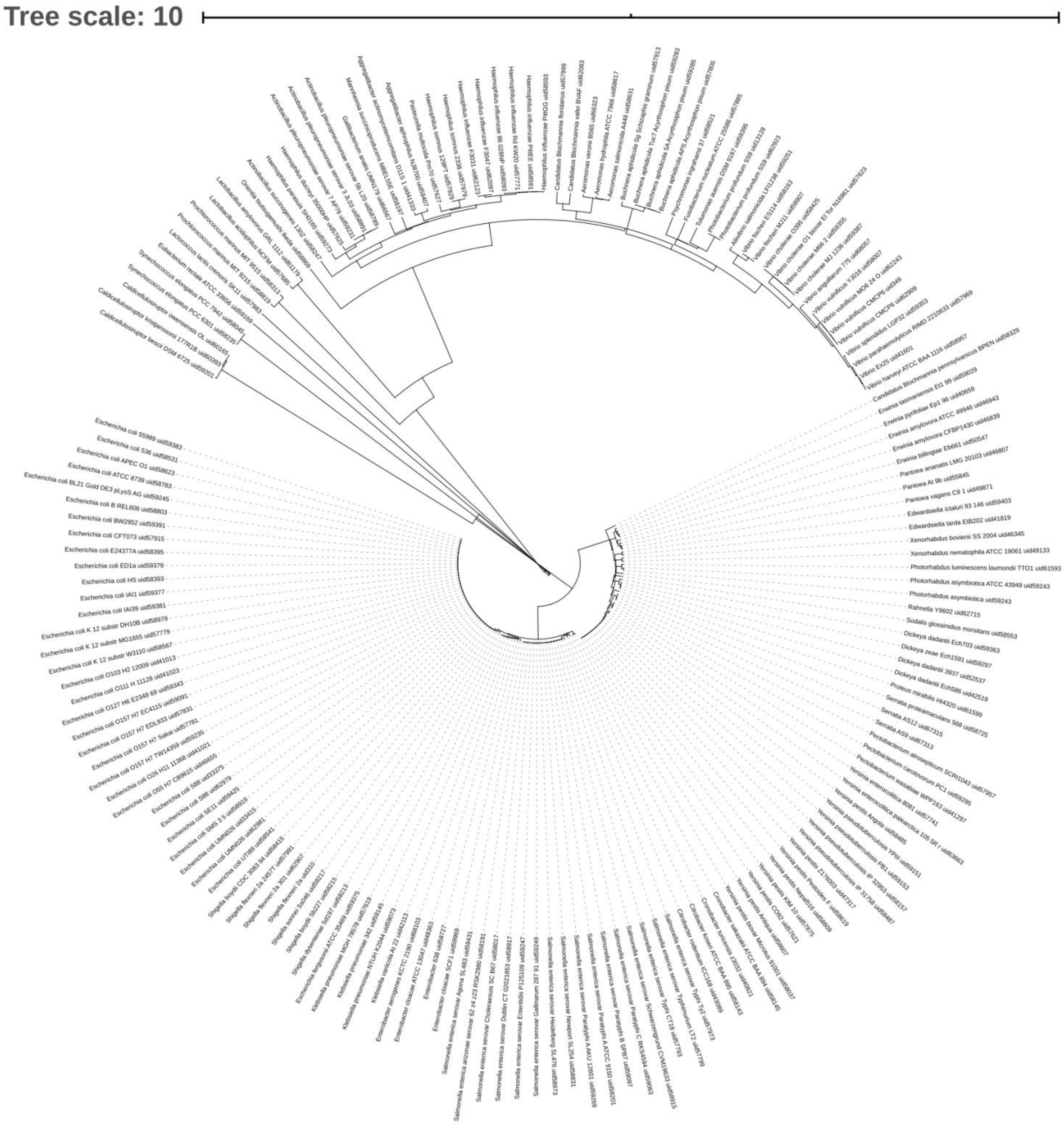


**Figure S1. The phylogenetic tree of TmaR**

A phylogenetic tree created by multiple sequence alignments of the amino acid sequence of TmaR (YeeX), as described in Material and Methods.

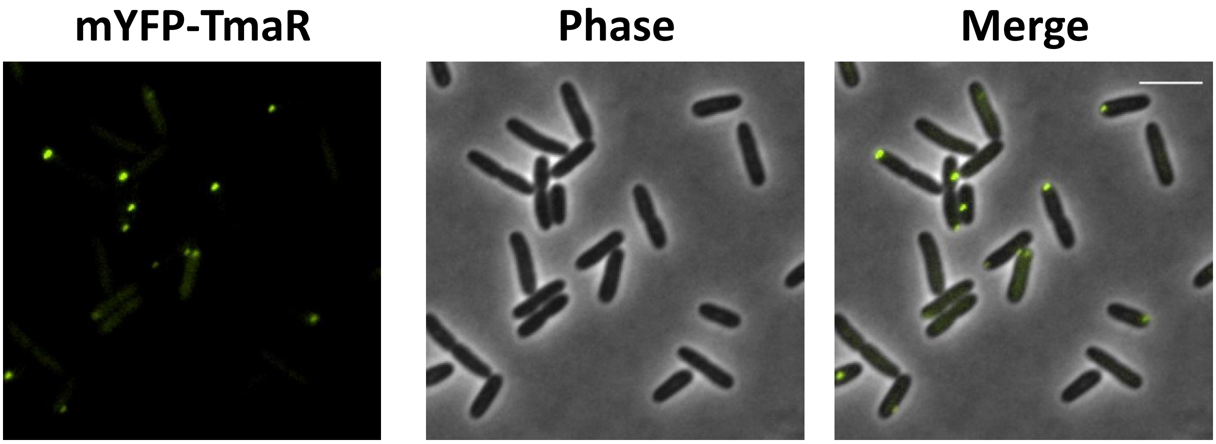


**Figure S2. mYFP-TmaR exhibits polar localization.**
Images of cells expressing TmaR, which is tagged at its N terminus with monomeric YFP (mYFP-TmaR).

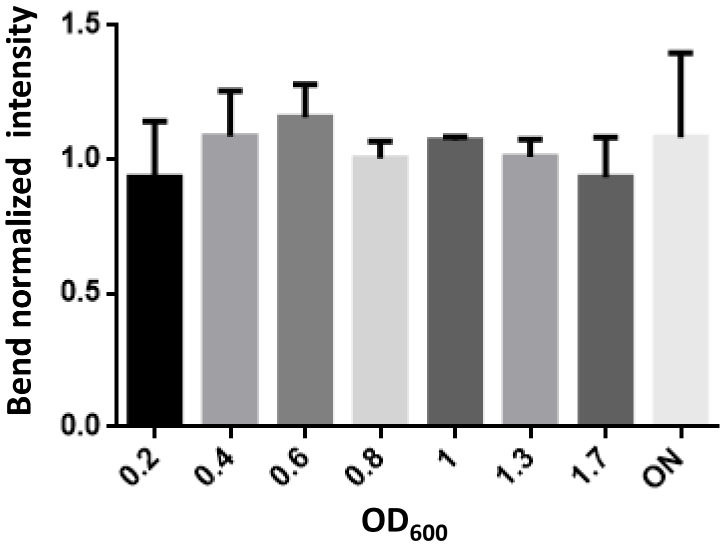


**Figure S3. The cellular level of TmaR in different growth phases is comparable.**
The average intensity of the TmaR-YFP bands in Fig. 1D normalized to the GroEL bands intensity in the same lysate. Cells were at the indicated $OD_{600}$ (ON, cells grown overnight). The bars show the standard error between three biological repeats.

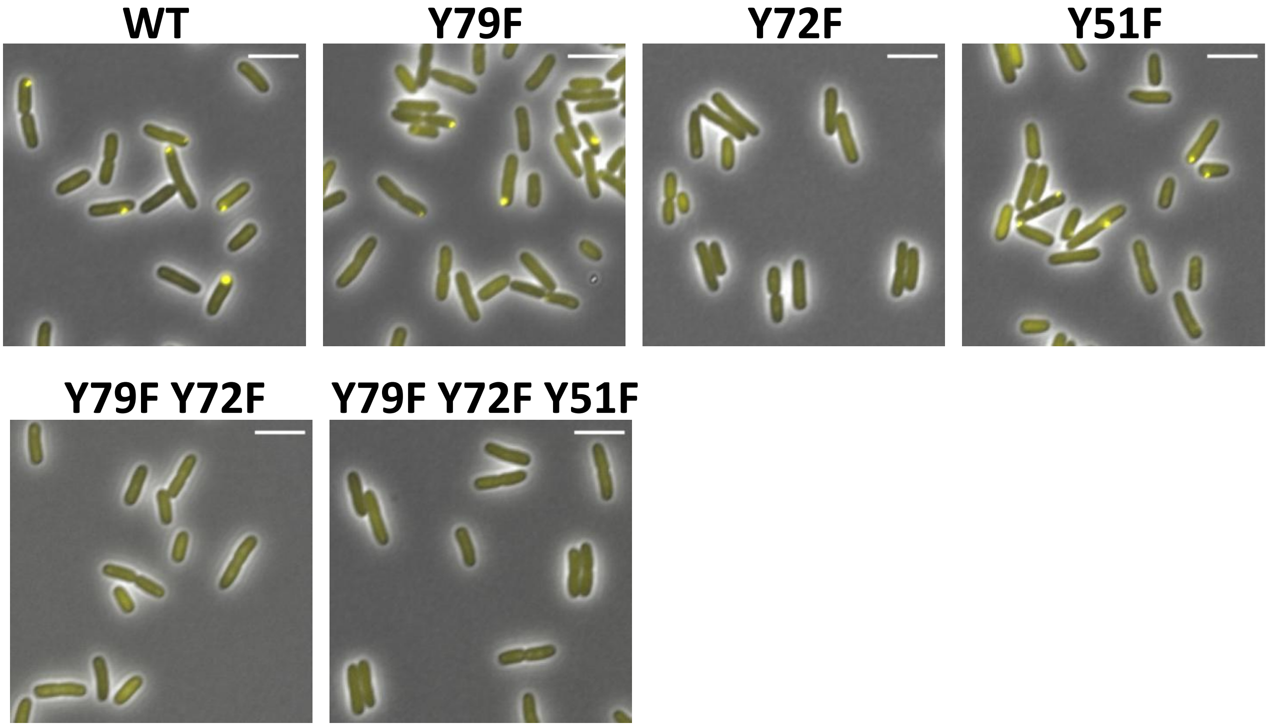


**Figure S4. Localization of mYFP-TmaR and its tyrosine replacements mutants.** Images of cells expressing mYFP-TmaR-YFP (WT) and its variants mutated in each of its three tyrosines, (Y79F, Y72F and Y51F), in two of its tyrosines (Y79F,Y72F) and in all three (Y79F,Y72F,Y51F), all tagged with mYFP.

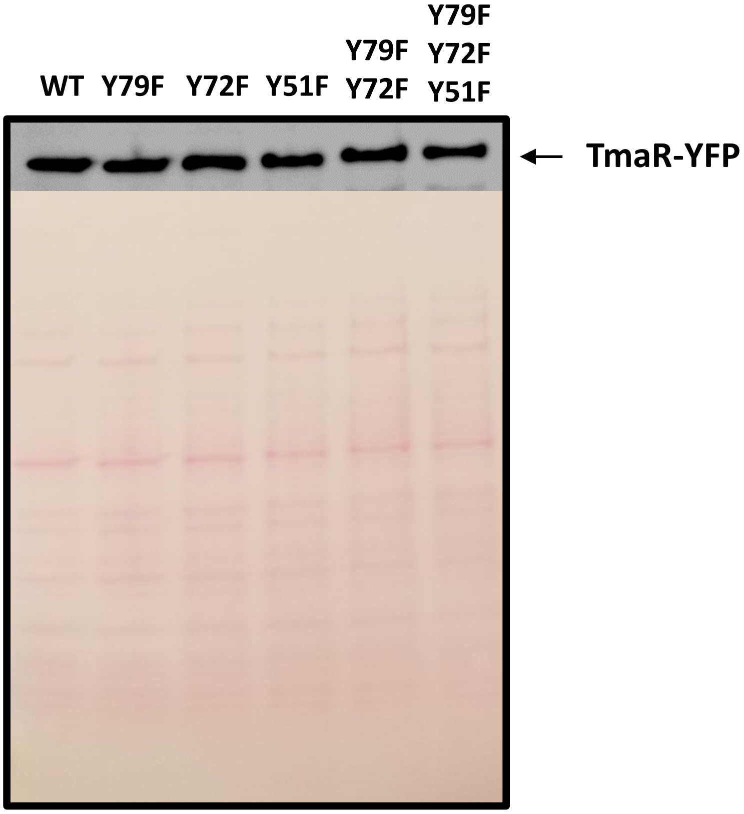


**Fig. S5. The mutations in TmaR tyrosines do not affect the level of TmaR**
Cells harboring plasmid-expressed mYFP-tagged TmaR (WT) or its variants mutated in each of the three tyrosines, (Y79F, Y72F and Y51F), in two of the tyrosines (Y79F,Y72F) and in all three tyrosines (Y79F,Y72F,Y51F), as well as cells with no plasmid (NC, negative control), were grown in LB to mid-logarithmic phase. The lysates were fractionated on a 4-20% SDS polyacrylamide gel and blotted onto membrane. The membrane was stained using Ponceau S (lower panel), to show that similar amount of lysates were loaded in each well, then washed and probed with anti-GFP2 (upper panel), to show that the substitutions of TmaR tyrosines did not change TmaR level.

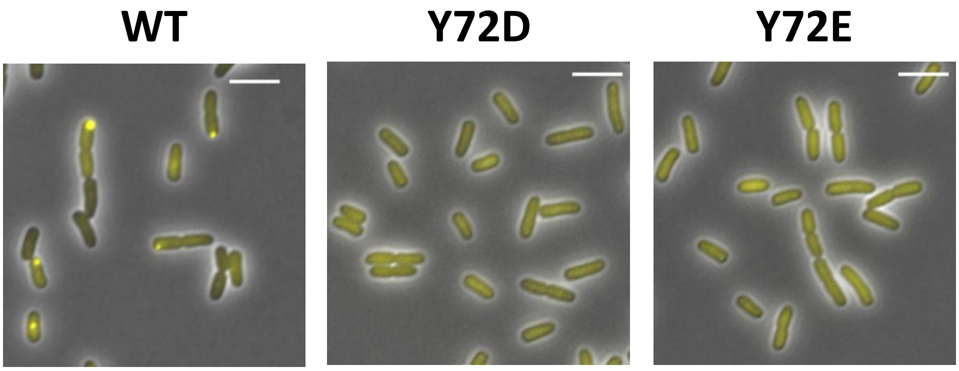


**Figure S6. Localization of both mYFP-TmaR Y72D and mYFP-TmaR Y72E is diffused.**
Images of cells expressing mYFP-TmaR (WT), mYFP-TmaR Y72D (Y72D) and mYFP-TmaR Y72E (Y72E).

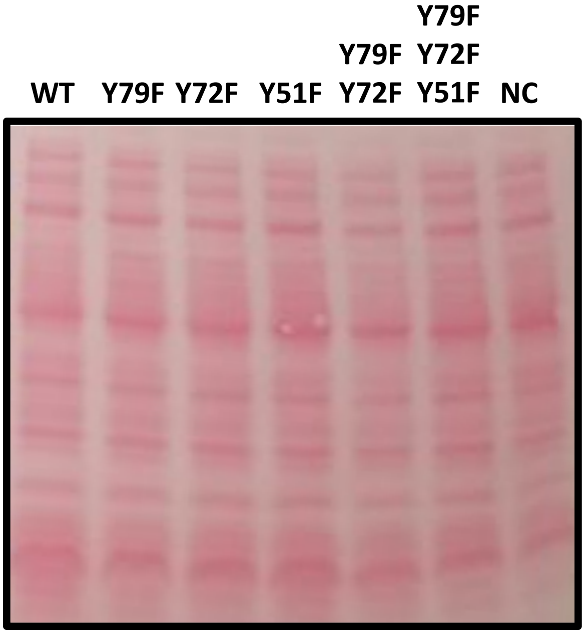


**Figure S7. Similar amounts of lysates were loaded to identify the phosphorylated tyrosine in TmaR by Western blot analysis** (supplemental information for Fig. 1G ).
For experimental details see the legend for Fig. 1G. The Ponceau S staining shows that a similar amount of lysates wereloaded in each well.

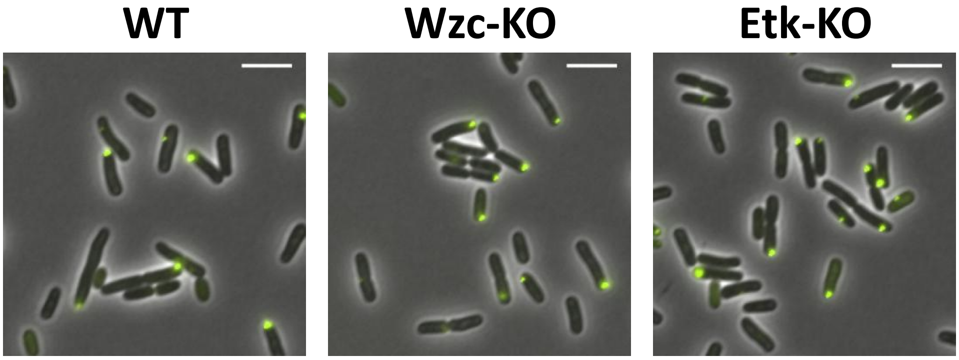


**Figure S8. Deletion of the known *E. coli* tyrosine kinases does not affect TmaR localization.**
Images of cells expressing TmaR-YFP in the background of *wzc* and *etk* genes (WT), in the background of *wzc* deletion (Wzc-KO) or in the background *etk* deletion (Etk-KO).

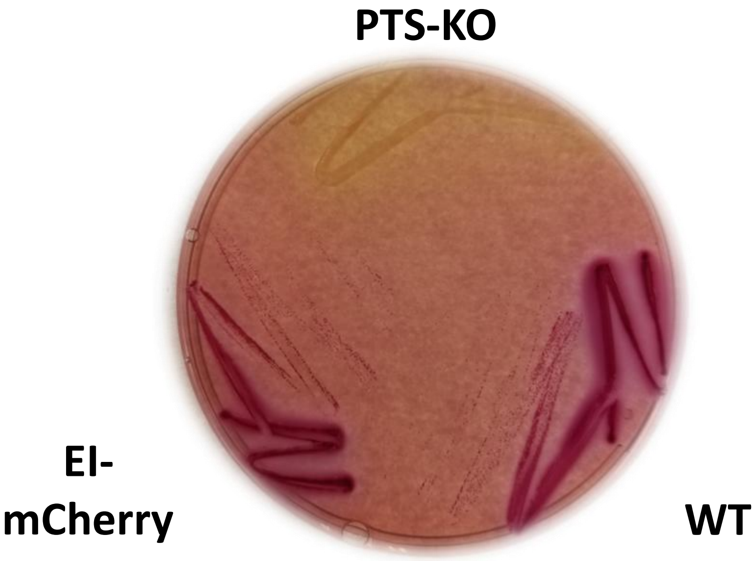


**Figure S9. EI-mCherry is active.**
A representative MacConkey plate showing the phenotype of cells expressing untagged EI (WT) or EI-mCherry, both expressed from EI endogenous promoter, as well as of cells deleted for the *pts* operon (PTS-KO).

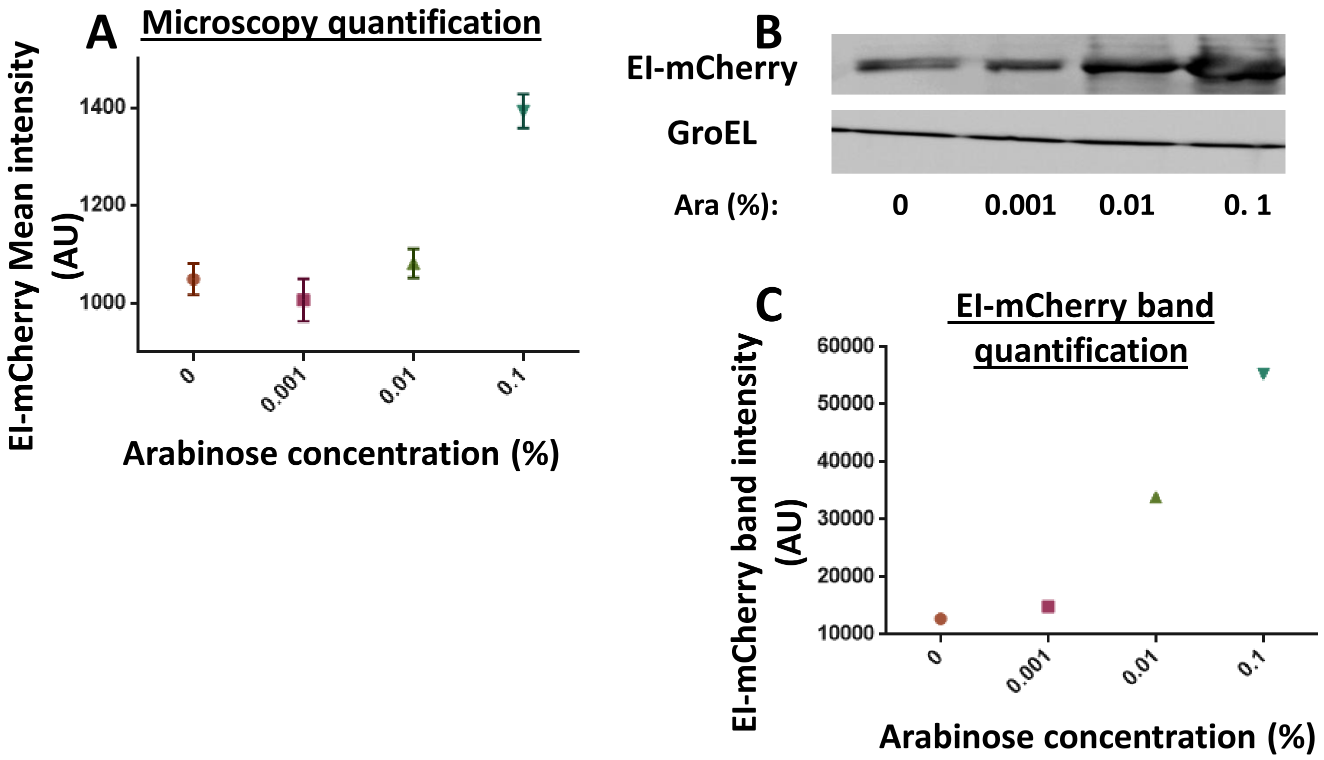


**Figure S10. EI-mCherry fluorescent signal correlates with EI-mCherry level.**
Cells expressing EI-mCherry from $P_{ara}$ in plasmid pBADLLEI-mCherry in the presence of different levels of the arabinose inducer until mid-logarithmic phase.
(A) The average of the mean EI-mCherry fluorescence upon induction by the indicated arabinose concentrations. The bars show the standard deviation between different fields.
(B) Western blot analysis showing EI-mCherry expression upon increasing concentration of arabinose, as indicated. Lysates were fractionated on a gel, blotted onto a membrane and probed with anti-mCherry antibodies (upper panel). The house-keeping gene GroEL served for normalizing the amounts loaded (lower panel).
(C) The average intensity of the EI-mCherry bands, normalized to the GroEL bands intensity, in the Western blot analysis presented in (B).

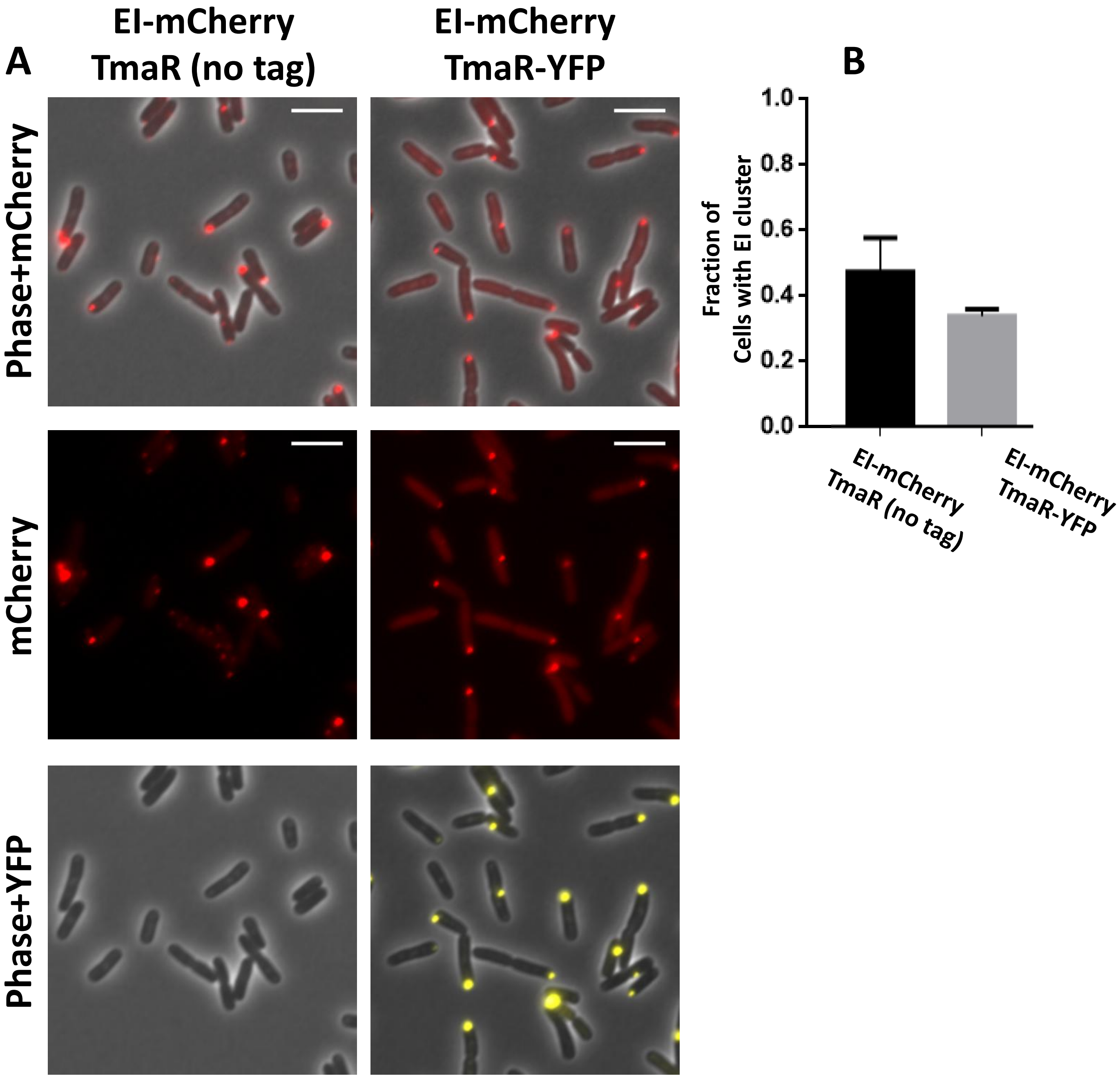


**Figure S11. Fluorescently-tagged and non-tagged TmaR have the same effect on EI-mCherry localization.**

(A) Images of cells expressing EI-mCherry in the presence of untagged TmaR (left) or YFP-tagged TmaR (right).

(B) The average fraction of cells with EI-mCherry clusters in the presence of untagged TmaR (black) and in the presence of TmaR-YFP (grey) in the experiment shown in (A). The bars show the standard deviation between different fields (N=200).

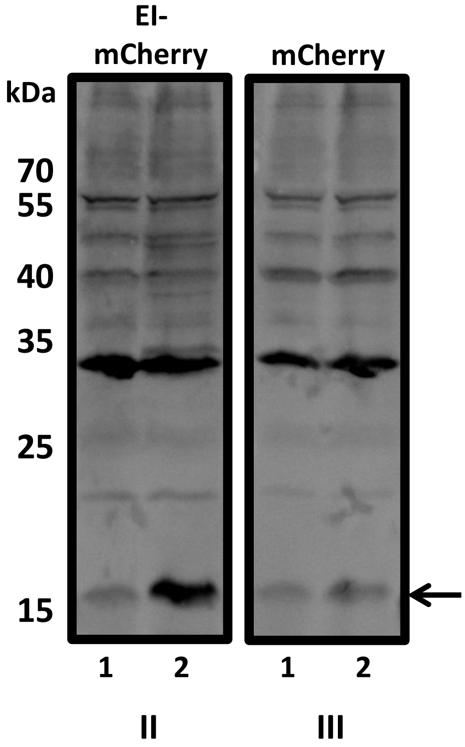


**Figure S12. Far Western analysis of the interaction between EI and TmaR.**
The full nitrocellulose membrane presented in Figure 2E. The following lysates were fractionated on the gel: Δ*tmaR* expressing the His-tag only (lanes 1) and Δ*tmaR* expressing His-tagged TmaR (lanes 2). The proteins were blotted onto a nitrocellulose membrane and probed with: EI-mCherry followed by anti-mCherry antibody (**II**) or mCherry followed by anti-mCherry antibody (**III**). Arrows point at the bands corresponding to His-TmaR size (~15 kDa).

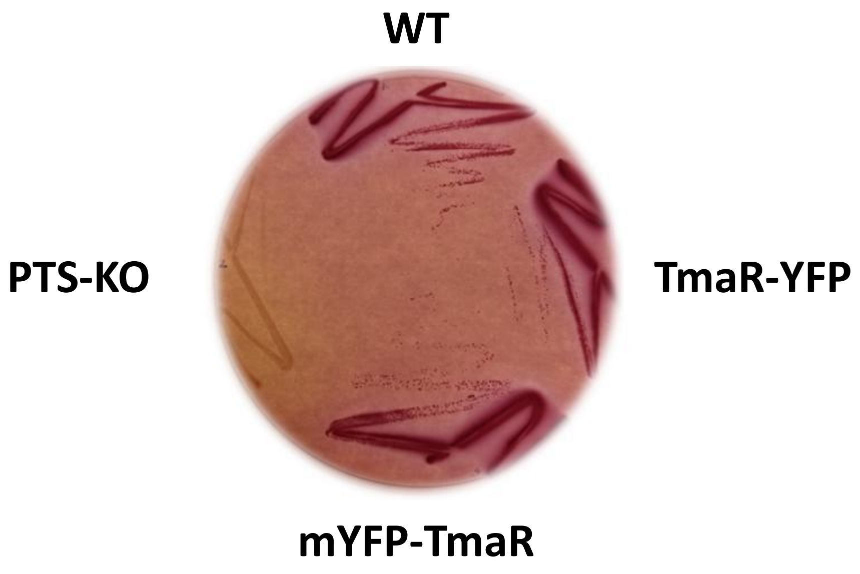


**Figure S13. EI expressed in the presence of TmaR tagged with YFP at its N- or C-terminus is as active as EI expressed in the presence of untagged TmaR.**
A representative MacConkey plate showing the phenotype of cells expressing untagged TmaR (WT), TmaR-YFP or mYFP-TmaR, all expressed from their endogenous promoter, as well as cells deleted for the *pts* operon (PTS-KO).

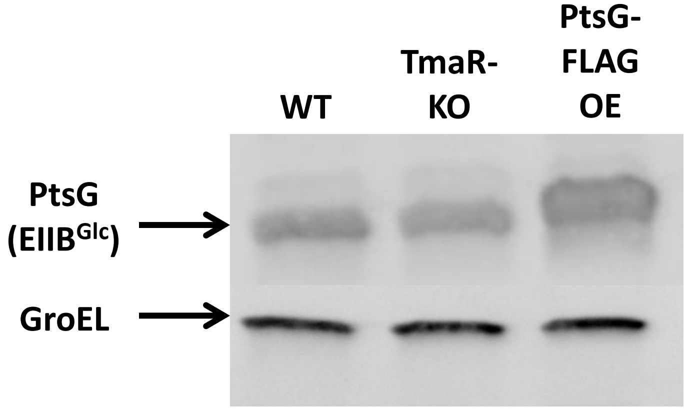


**Figure S14. Cells expressing TmaR or deleted for it show the same level of PtsG ($IIECB^{Glc}$).**
Western blot analysis of lysates of wild-type cells (WT), cells deleted for the *tmaR* gene (TmaR-KO) and cells overexpressing PtsG-FLAG (PtsG OE), which were fractionated on a gel, blotted onto a membrane and probed with anti-$IIB^{Glc}$ antibodies. The house keeping gene GroEL served for normalization.

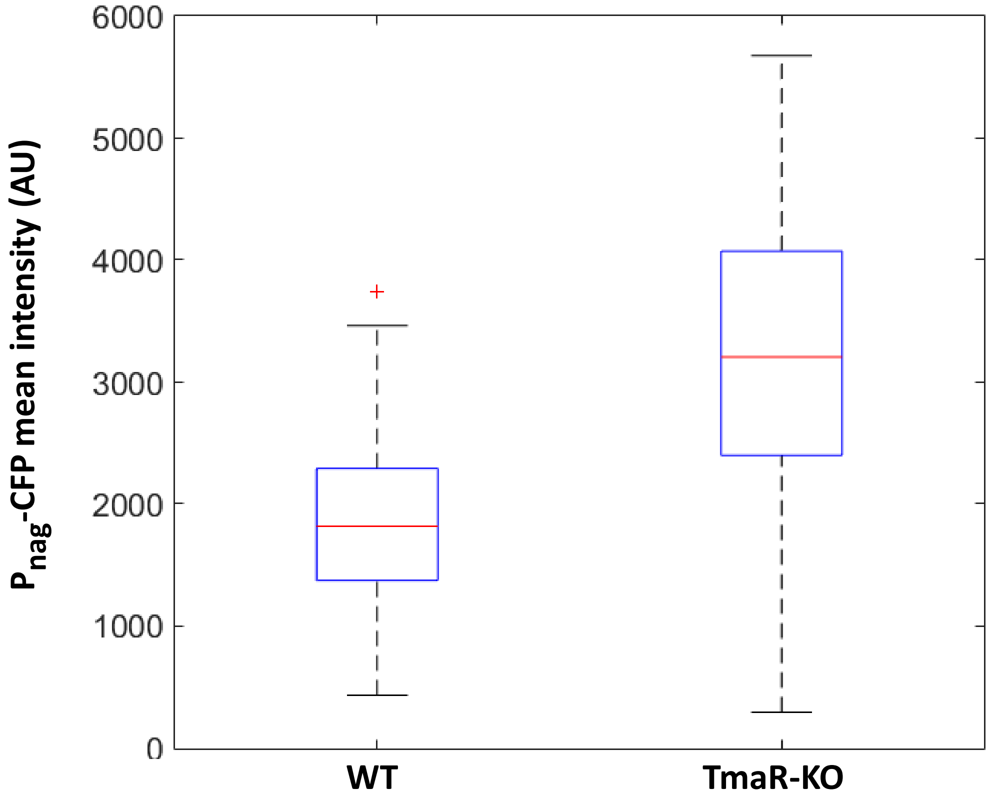


**Figure S15. The TmaR-KO cells show higher heterogeneity in PTS sugar consumption than WT cells.**

A box plot showing the mean intensity (arbitrary units, AU) of mCerulean expressed from $P_{NAG}$ in WT and TmaR-KO cells 1 hour after their transition to NAG-containing medium. The central mark in each box indicates the median, and the bottom and top edges of the boxes indicate the 25$^{th}$ and 75$^{th}$ percentiles, respectively. The whiskers extend to the most extreme data points, which are not considered outliers, and the outliers are plotted individually using the '+' symbol.

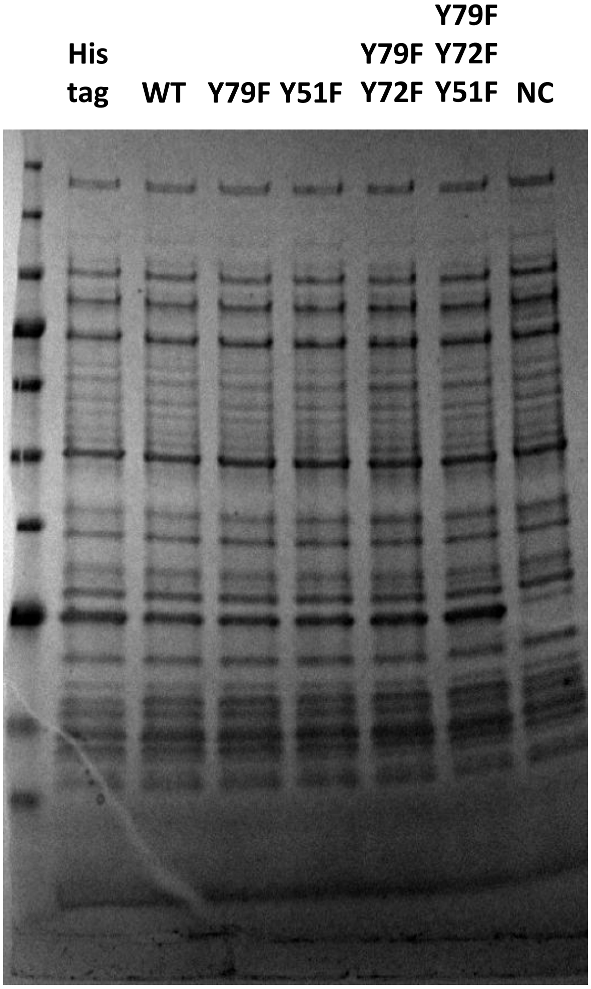


**Figure S16. Equal amounts of lysates were loaded for the Far Western analysis of EI interaction with TmaR pTyr mutants (**supplemental information for Fig. 5A **).**
Lysates of Δ*tmaR* cells overexpressing from his tag only or TmaR (WT) and its variants mutated in each of its three tyrosines, (Y79F and Y51F), in two of its tyrosines (Y79F,Y72F) and in all three (Y79F,Y72F,Y51F), as well as Δ*tmaR* cells with no plasmid (NC, negative control) were fractionated on a gel. The gel was stained by Coomassie Brilliant Blue. The arrow points at TmaR proteins (~15 kDa).

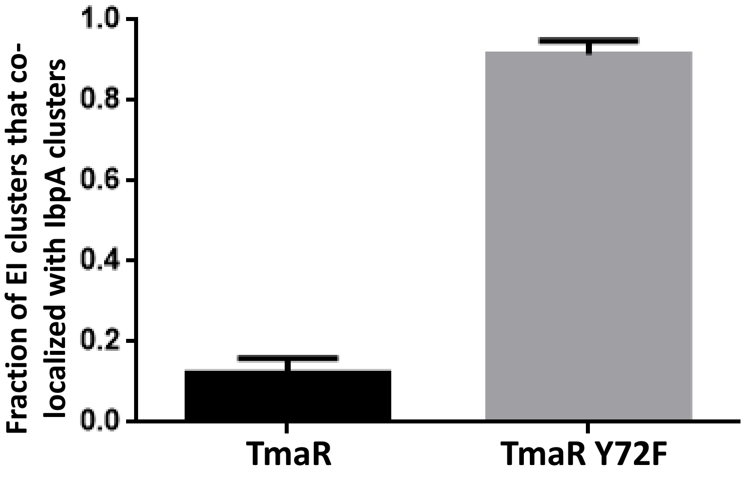


**Figure S17. EI aggregates to form inclusion bodies only in the presence of non-phosphorylate TmaR (**supplemental information for Fig. 5A **).**
The average fraction of EI-mCherry clusters that co-localized with IbpA-msfGFP clusters in cells overexpressing non-tagged TmaR or TmaR Y72F. The bars show the standard deviation between different fields (N=150).

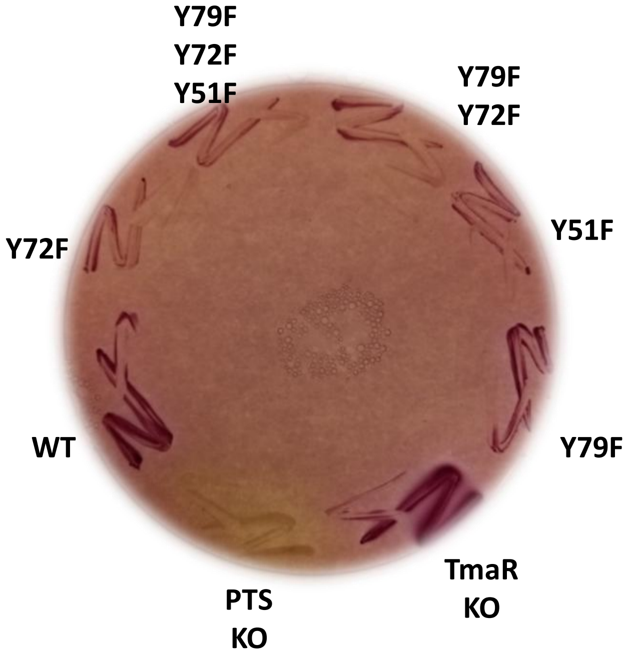


**Figure S18. Cell expressing all TmaR mutants with tyrosine substitutions are capable of consuming sugar to a certain extent**
A representative MacConkey plate showing the phenotype of Δ*tmaR* cells , WT cells and cells expressing TmaR mutants with the indicated substitutions from *tmaR* native promoter and locus in the chromosome, as well as cells deleted for the *pts* operon (PTS-KO), which served as a negative control for sugar consumption.

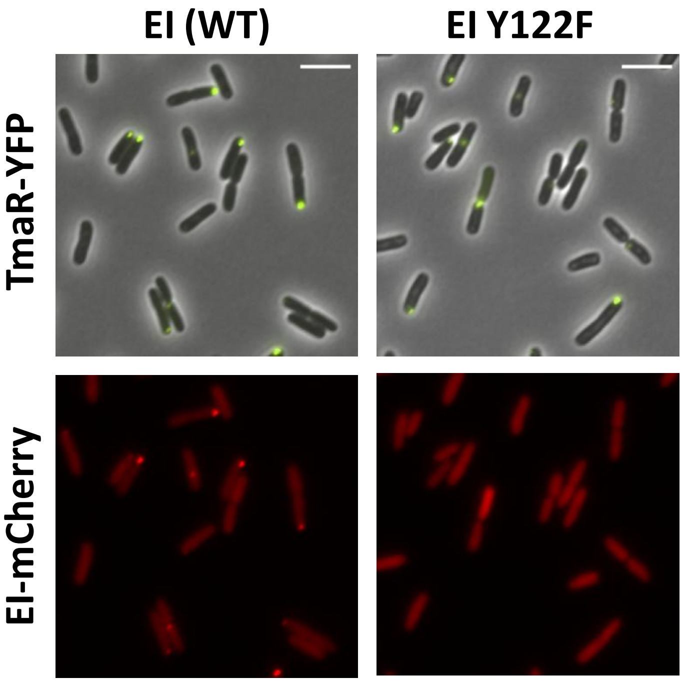


**Figure S19. The EI Y122F mutant protein does not affect TmaR localization**
Images of cells co-expressing TmaR-YFP with either EI-mCherry (WT, left panels) or EI Y122F-mCherry (EI Y122F, right panels). TmaR-YFP is in yellow and mCherry-tagged EI proteins (WT and Y122F) are in red.

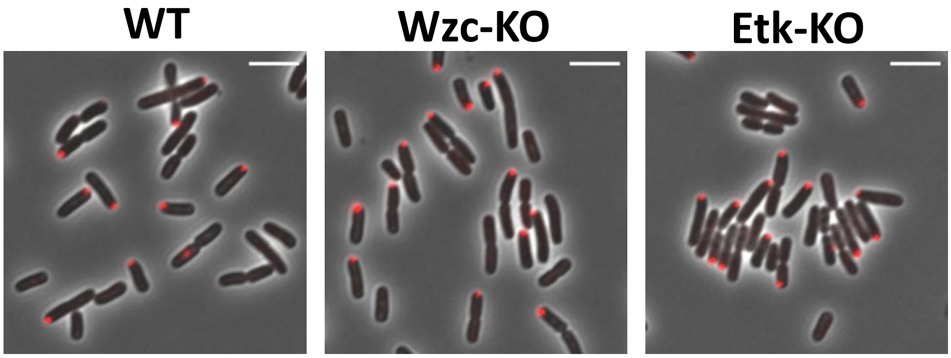


**Figure S20. Deletion of the known *E. coli* tyrosine kinases does not affect EI localization**
Images of cells expressing EI-mCherry in the background of *wzc* and *etk* genes (WT), in the background of *wzc* deletion (Wzc-KO) or in the background *etk* deletion (Etk-KO).

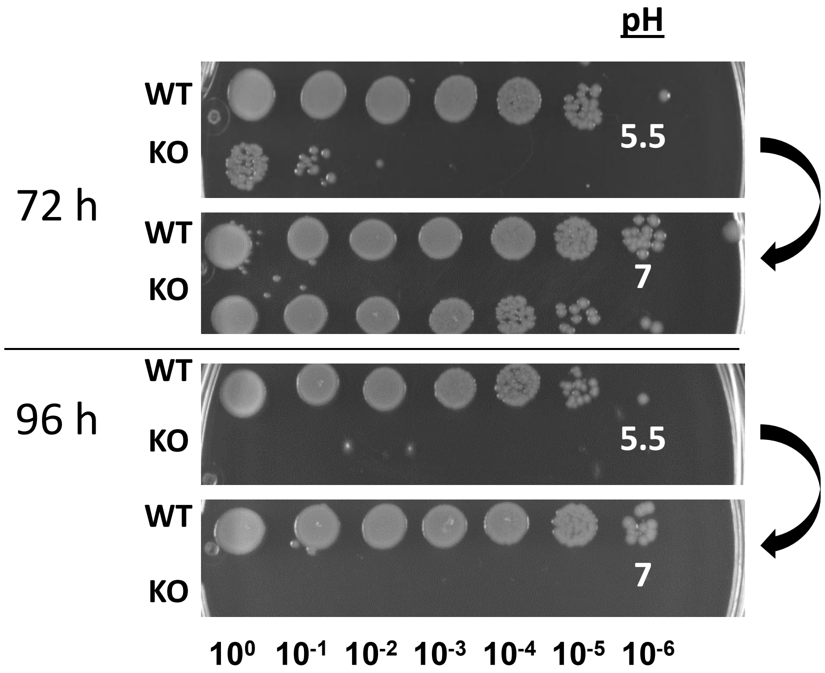


**Figure S21. The drop in survival of TmaR-KO in acidic pH is reversible only after 72h, but not after 96h**
Pictures of wild-type (WT) and TmaR-KO (KO) colonies generated by cells grown in acidic M9 medium (pH 5.5) supplemented 0.4% glucose for 72 or 96 hrs (as indicated), which were diluted 1:100 into M9 (pH 7) medium and kept at regular growing conditions for additional 24 hrs.

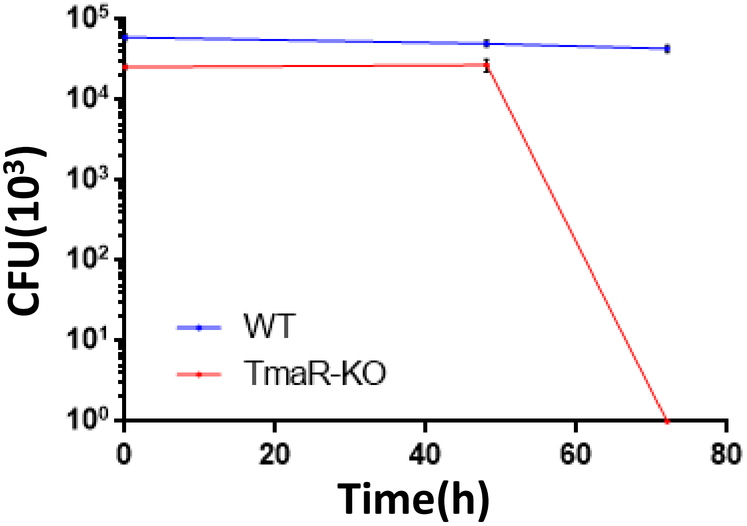


**Figure S22. The reduction in survival of cells lacking TmaR in mild acidic conditions occurs between 48 and 72 hours of growth.**
The number of colonies formed (CFU), when plating 1 ml of wild type (WT, blue) or Δ*tmaR* (TmaR-KO, red) cells that were grown in acidic M9 medium (pH 5.5) supplemented with 0.4% glucose at pH 5.5, after 1, 2 and 3 days. The bars show the standard deviation between the two biological repeats.

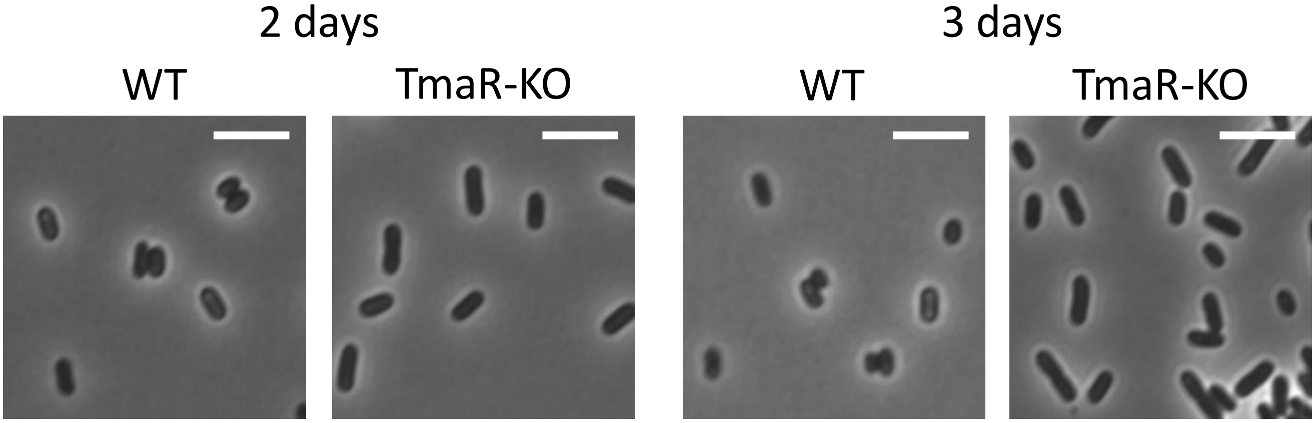


**Figure S23. TmaR-KO cells exhibit different morphology then TmaR WT cells after two and three days**

Images of WT or TmaR-KO cells that were picked from a MacConkey plate after two (right panel) or three (left panel) days.

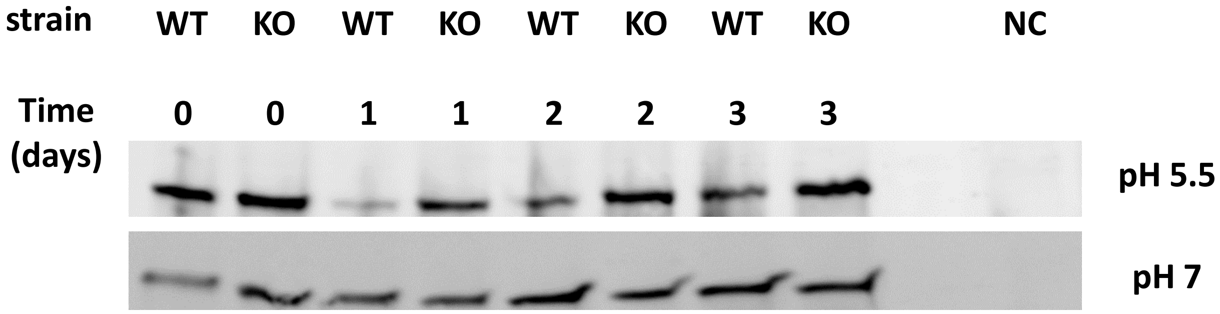


**Figure S24. Western blot analysis showing that RpoS level drops in wild-type cells after one day of growth, but not in TmaR-KO cells, in acidic, but not in neutral medium**

Equal amounts of lysates of WT, TmaR-KO (KO) and *ΔrpoS (*negative control, NC) cells grown in acidic (pH 5.5) or neutral (pH 7) medium supplemented with 0.4% glucose for 1, 2 or 3 days (as indicated) were fractionated on a 4-20% gradient SDS polyacrylamide gel, blotted onto a nitrocellulose membrane and probed with α-RpoS antibodies.

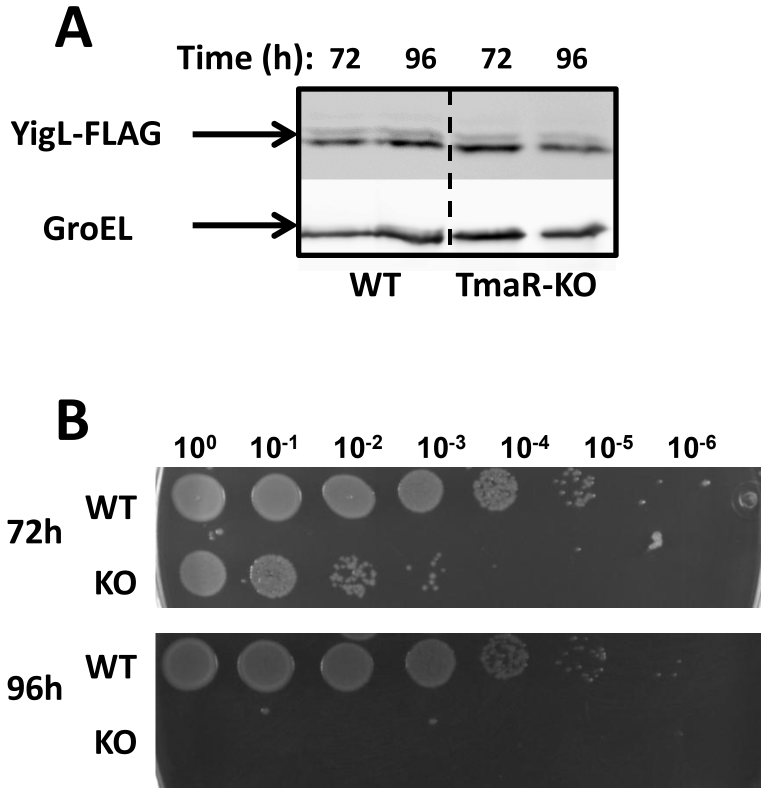


**Figure S25. TmaR-KO cells do not show an increase in YigL expression level**
(A) Western blot analysis of wild-type (WT) and Δ*tmaR* (TmaR-KO) cells, both expressing FLAG-tagged YigL from *yigL* native promoter and locus in the chromosome. Cells were grown in acidic M9 medium (pH 5.5) supplemented with 0.4% glucose at pH 5.5 for 72h or 96h (as indicated). Cell lysates were fractionated on a 12% SDS polyacrylamide gel, blotted onto a nitrocellulose membrane and probed with anti-FLAG antibodies. The house keeping gene GroEL served for normalization.
(B) Pictures of wild-type (WT) and Δ*tmaR* (KO) colonies expressing YigL-FLAG grown in acidic M9 medium (pH 5.5) supplemented with 0.4% glucose for 72h or 96h (as indicated).

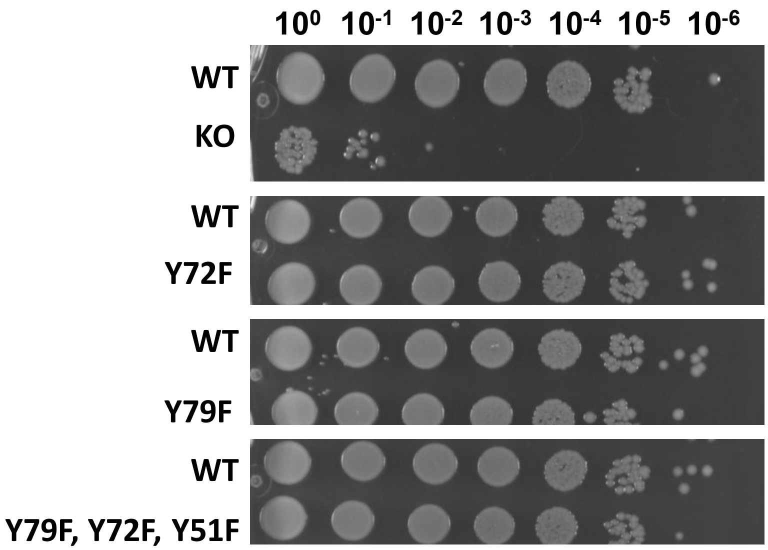


**Figure S26. The effect of mutations that prevent of TmaR phosphorylation on survival in acidic medium**

Pictures of wild-type (WT), TmaR-KO (KO) and TmaR mutants ( Y72F, Y79F and Y79F Y72F Y51F) expressed from TmaR native promoter. The cells were grown in acidic medium (pH 5.5) supplemented with 0.4% glucose for 72 hours.
